# Innate immune epigenomic landscape following controlled human influenza virus infection

**DOI:** 10.1101/2024.09.20.612974

**Authors:** William Thistlethwaite, Sindhu Vangeti, Wan-Sze Cheng, Pankaj Agarwal, Antonio Cappuccio, Wenliang Wang, Bei Wei, Rachel Myers, Aliza B. Rubenstein, Daniel Chawla, Manoj Hariharan, Micah T. McClain, Thomas W. Burke, Steven H. Kleinstein, Joseph R. Ecker, Christopher W. Woods, William J. Greenleaf, Xi Chen, Irene Ramos, Elena Zaslavsky, Thomas G. Evans, Olga G. Troyanskaya, Stuart C. Sealfon

## Abstract

Viral infections can induce changes in innate immunity that persist after virus clearance. Here, we used blood samples from a human influenza H3N2 challenge study to perform comprehensive multi-omic analyses. We detected remodeling of immune programs in innate immune cells after resolution of the infection that was proportional in magnitude to the level of prior viral load. We found changes associated with suppressed inflammation including decreased cytokine and AP-1 gene expression as well as decreased accessibility at AP-1 targets and interleukin-related gene promoter regions. We also found decreased histone deacetylase gene expression, increased MAP kinase gene expression, and increased accessibility at interferon-related gene promoter regions. Genes involved in inflammation and epigenetic-remodeling showed modulation of gene-chromatin site regulatory circuit activity. These results reveal a coordinated rewiring of the epigenetic landscape in innate immune cells induced by mild influenza virus infection.

## Introduction

While vertebrate immunity has been classically separated into an innate immune system that lacks immunological memory and an adaptive immune system that targets specific pathogens, there is growing awareness of “innate immune memory,” encoded by epigenetic changes in innate immune cells induced by perturbations such as infection or vaccination.^1,2^ These epigenetic changes, including histone modifications, DNA methylation, and chromatin remodeling, can persist after the acute phase of the infection and can broadly alter the response to a subsequent immune challenge unrelated to the earlier perturbation.^3^

In some instances, viral infections cause epigenetic changes that result in prolonged dysregulation of the innate immune system. Severe COVID-19 in humans caused persistent myeloid dysregulation, marked by long-lasting inflammation and immunosuppression in the neutrophil compartment.^4^ Asymptomatic and mild SARS-CoV-2 infection caused prolonged pro-inflammatory changes in DNA methylation sites associated with poorer responses to infection and resembling signatures associated with chronic infections and autoimmune diseases.^5^ Persistent methylation changes associated with airway inflammatory processes have been found following respiratory syncytial virus (RSV) infections.^6^ However, epigenetic changes induced by viral infections have also been shown to augment responses to secondary infections and to vaccinations.^7,8^

While influenza virus infection is known to induce changes in cross-reactive immunity (for example, to bacterial infection),^9,10^ *in vivo* studies of the effects of influenza infection on humans have largely focused on acute responses and transcriptome analyses.^3^ *In vitro* studies of A549 cells have shed some light on how the influenza virus may shape the epigenetic landscape,^11,12^ but these studies do not capture the dynamic nature of the innate immune system or whether epigenetic changes persist after the acute infection is cleared. While prolonged epigenetic remodeling following influenza vaccination has been reported,^13^ little is known about innate immune epigenetic remodeling following the resolution of acute influenza virus infection.

Here, we performed a multiomic integrated analysis in a human influenza H3N2 challenge study. We investigated whether innate immune cell subtypes developed different epigenetic profiles after resolution of this mild infection at 28 days post-challenge. A controlled human infection provided a unique opportunity to characterize precisely timed molecular changes relative to pre-infection state. We conducted single-cell and bulk sample multiomic analyses of chromatin accessibility, transcriptome, DNA methylation, and histone modification assays to provide a high resolution characterization of the coordinated changes in the epigenetic memory landscape following infection.

## Results

### Study design and data generation

Data were generated using cryopreserved PBMCs and PaxGene blood RNA samples collected from the challenge phase of a universal influenza A vaccine clinical trial.^14^ In the vaccination phase of the trial, 45 subjects randomized to mock-vaccination with 0.5 ml of a saline placebo (“naive”) and 69 subjects randomized to vaccination with 0.5 ml of a 1.5×10^8^ pfu (4.3 ×10^8^ TCID_50_/ml) dose of MVA-NP+M1 vaccine (“vaccinated”).

Approximately six weeks after saline or vaccine administration, all study subjects were challenged intranasally with 0.5 ml of a 1×10^6^ TCID_50_/ml dose of influenza A (H3N2) virus. All subjects tested seronegative to the challenge strain prior to inclusion. A schematic of the challenge phase, including data generation for our study, is shown in **Figure 1A**. As previously reported, vaccination induced significant increases in CD4+ and CD8+ T cell response, but did not impact viral load or clinical outcome after challenge.^15^ In order to capture the regulatory responses to the infection without any confounds introduced by the recent vaccination, our primary analyses focused on the vaccine-naive subjects, and data obtained from the virus challenge samples in vaccinated subjects were used for comparison purposes. Demographics are included in **Table S1** (naive group) and **Table S2** (vaccinated group).

**Figure 1.**
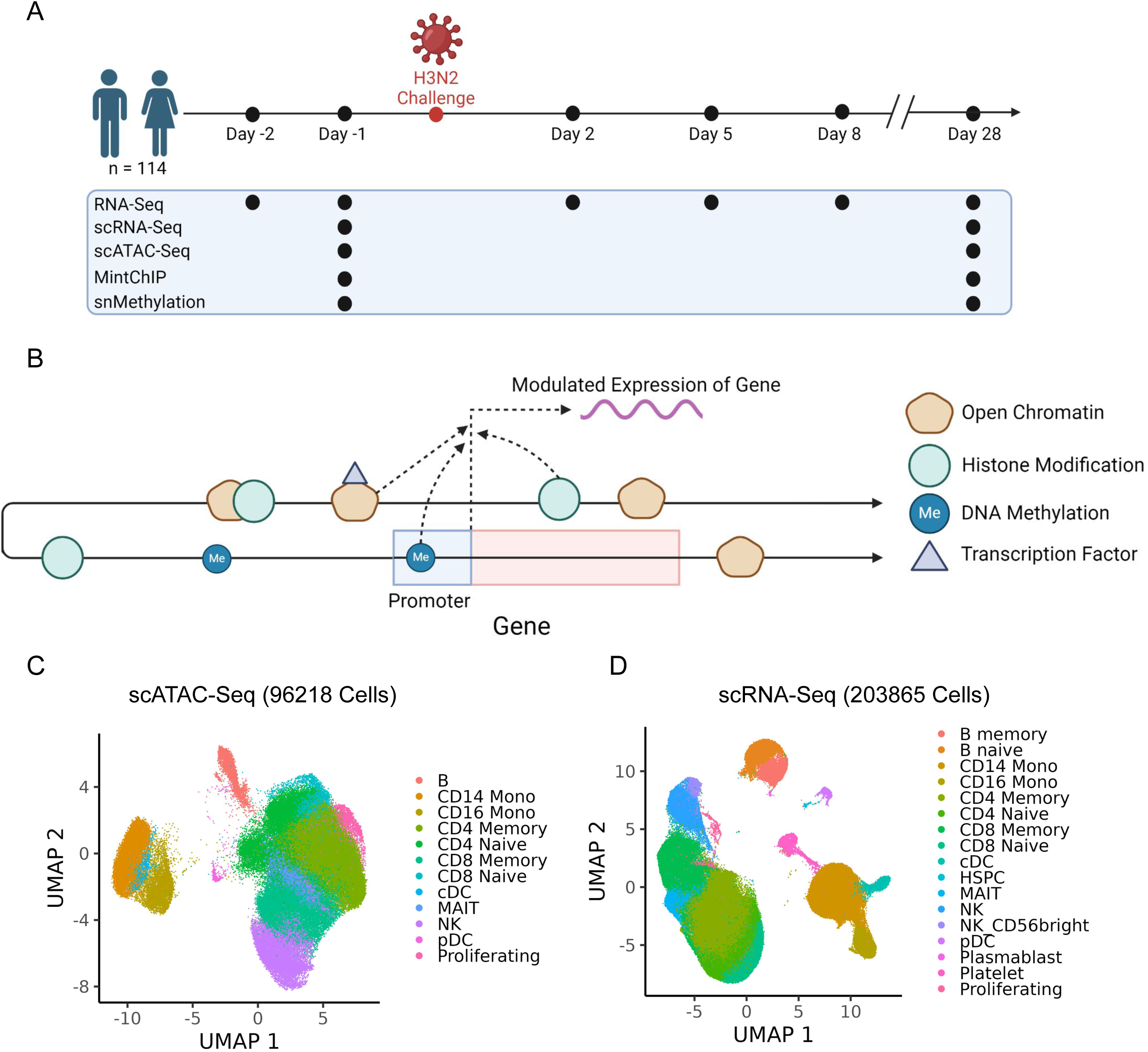
Overview of human influenza H3N2 challenge study. (See also Tables S1-S2.) (A) Schematic depicting time course of human influenza H3N2 challenge study. A total of 114 subjects were challenged intranasally with influenza A (H3N2) virus. Subjects were epigenetically profiled 1 day pre-challenge and 28 days post-challenge and were transcriptionally profiled throughout the entire time course. (B) Schematic of different epigenetic mechanisms captured in the study that can modulate gene expression. (C) Overview of scRNA-seq data at 1 day pre-challenge and 28 days post-challenge. Data come from 10 subjects (20 samples in total). (D) Overview of scATAC-seq data at 1 day pre-challenge and 28 days post-challenge. Data come from 10 subjects (20 samples in total).

Gene expression is controlled in part by the rate of transcription, which is modified by epigenetic mechanisms including histone modification, DNA methylation, and changes in chromatin accessibility occurring at proximal (promoter) and distal (enhancer, repressor) regulatory sites (**Figure 1B**). Regulatory sites that are not adjacent on the linear DNA sequence can interact in 3D space to recruit transcription factors and other regulatory factors that modulate gene transcription and RNA levels. To elucidate these coordinated processes, subjects were studied with complementary molecular assays. Subjects were divided into three groups based on their real-time PCR estimated viral loads following challenge (see Methods). The viral load groups comprised low viral load (LVL, n = 16 for naive and n = 22 for vaccinated), moderate viral load (MVL, n = 10 for naive and n = 25 for vaccinated), and high viral load (HVL, n = 19 for naive and n = 22 for vaccinated). Single-cell chromatin accessibility data (profiled with scATAC-seq, **Figure 1C**) and transcriptome data (profiled with scRNA-seq, **Figure 1D**) were obtained from a total of three LVL and seven HVL subjects. Thirteen naive HVL subjects were profiled by single-nucleus DNA methylation^16^ and eleven naive HVL subjects were profiled by multiplexed, indexed T7 ChIP-seq (Mint-ChIP)^17^ to measure histone levels for 6 different histone features. Thus, naive HVL subjects were profiled more deeply than other subjects and were the primary focus of analyses below. In addition, samples from four vaccinated subjects were selected for single-cell chromatin accessibility (scATAC-seq) and transcriptome (scRNA-seq) profiling (see Methods). Finally, whole blood samples from all subjects were profiled with bulk transcriptomics (RNA-seq) at all time points to connect changes seen in the post-infection multiome data to transcription during the acute phase of infection.

### Mild influenza virus infection induces persistent epigenetic and transcriptional changes in the innate immune system

In order to evaluate the coordinated molecular responses present after resolution of the acute infection, we first studied differences between pre-challenge and 28 day post-challenge samples in naive HVL subjects. scATAC-seq identified 7,426 differentially accessible chromatin sites (DAS; **Figure 2A** and **Table S3**) across 5 innate immune cell types. The genomic locations of the DAS in naive HVL subjects were concentrated in promoter regions in all cell types (**Figure 2B**). scRNA-seq identified 621 differentially expressed genes (DEG; **Figure 2C** and **Table S4**) across 6 innate immune cell types. No statistically significant cell type proportion changes between pre- and 28 day post-challenge samples were observed. Next, we assessed if the molecular responses established in HVL subjects followed similar patterns to those found in LVL subjects (**Table S5** and **Table S6**). The fold change magnitude of the DEG and DAS present 28 days after challenge were in general higher in the HVL than in the LVL subjects, with the direction of change significantly correlated in pDCs, CD16 monocytes, cDCs, NK cells, and CD14 monocytes for DEG (**Figure 2D**) and in NK cells and CD14 monocytes for DAS (**Figure S1A**). When comparing the virus challenge single-cell assay results in naive subjects to those in vaccinated subjects (**Table S7** and **Table S8**), DEG fold changes (FC) were significantly correlated for all major innate immune cell types (**Figure S1B**) while DAS FC correlation achieved significance in CD14 monocytes (**Figure S1C**). The poorer correlation seen in DAS relative to DEG most likely results from the scATAC-seq data being much sparser than the scRNA-seq data. Because the majority of naive HVL subjects were female and all of the vaccinated HVL subjects were male, the similarity of the single-cell responses to challenge for the naive and vaccinated subjects also suggests that many of the transcriptomic changes observed after resolution of initial infection are shared across the sexes.

**Figure 2.**
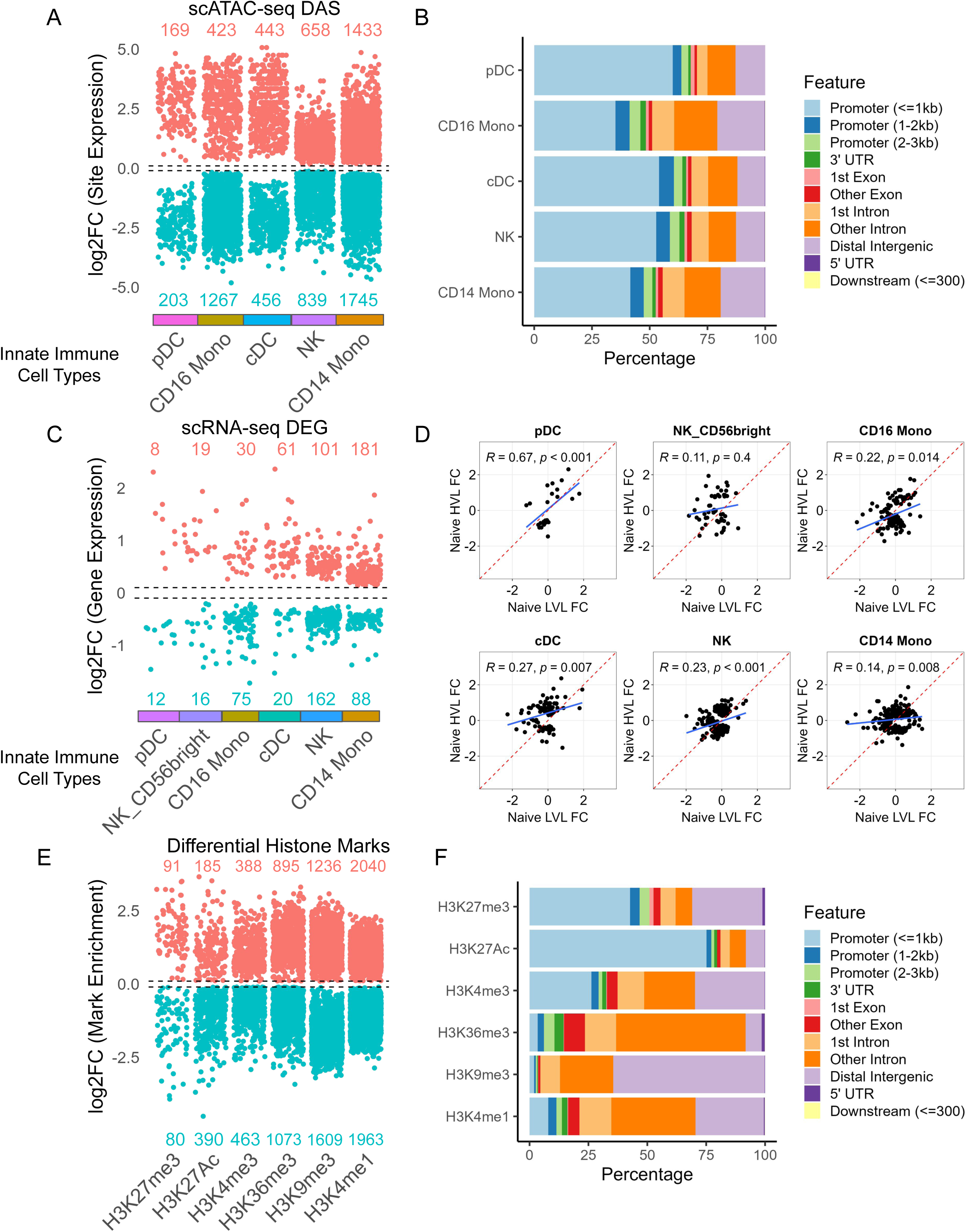
Mild influenza virus infection induces long-lasting epigenetic and transcriptional changes. (See also Figure S1 and Tables S3-S9.) All differential expression analyses compared data from 28 days post-challenge to 1 day pre-challenge. (A) Differentially accessible sites (DAS) found in scATAC-seq data for innate immune cell types. Fold change threshold of 0.1 was used. The number of upregulated and downregulated DAS are listed for each cell type. Cell type color bars match the scATAC-seq UMAP cell type colors. (B) Distribution of genomic features for scATAC-seq DAS. (C) Differentially expressed genes (DEG) found in scRNA-seq data for innate immune cell types. Fold change threshold of 0.1 was used. The number of upregulated and downregulated DEG are listed for each cell type. Cell type color bars match the scRNA-seq UMAP cell type colors. (D) For each scRNA-seq cell type, fold change correlations (Spearman) between naive high viral load (HVL) subjects and naive low viral load (LVL) subjects were calculated for the union of naive HVL and LVL DEGs (28 days post-challenge versus pre-challenge). (E) Differentially enriched histone marks found in Mint-ChIP data. Fold change threshold of 0.1 was used. The number of upregulated and downregulated sites are listed for each histone mark. (F) Distribution of genomic features for differentially enriched histone marks.

Histone modification analysis by Mint-ChIP comparing pre- to 28 day post-challenge in naive HVL subjects identified 10,413 regulated loci across all marks assayed (H3K27ac, H3K4me1, H3K4me3, H3K9me3, H3K27me3, and H3K36me3; **Figure 2E** and **Table S9**). The genomic locations for regulated loci for each histone mark were consonant with expectations (e.g., H3K27ac was enriched in promoter regions and H3K36me3 was enriched in gene bodies; **Figure 2F**).^18,19^ The majority of enriched loci for histone mark H3K27ac, which is associated with active transcription, were downregulated. As previously reported as part of a snMethylation survey of regulatory responses across a large number of conditions,^16^ analysis of 7 sorted immune cell types from the influenza challenge study revealed more than 70,000 differentially methylated regions (DMR) between pre- and post-challenge samples in naive HVL subjects. Overall, these analyses revealed extensive epigenetic and transcriptional changes were present in innate immune cells after clearing the acute infection.

### Analysis of differentially accessible sites reveals regulatory programs related to inflammation and epigenetic remodeling

We next characterized the biological processes represented by the differential regulatory features observed at 28 days after infection in innate immune cells, focusing first on the genes showing differential promoter accessibility in the naive HVL scATAC-seq data. Functional module discovery analysis^20^ identified upregulated (Figure 3) and downregulated (Figure S2) modules related to inflammation and epigenetic remodeling in CD14 and CD16 monocytes as well as conventional dendritic cells (cDCs). Genes showing increased accessibility were annotated to MAP kinase processes, including positive regulation of MAPK cascade, ERK1 and ERK2 cascade, apoptotic signaling, and programmed cell death, as well as response to interferon-gamma and interferon-beta. Downregulated DAS were associated with downregulation of cell adhesion, interleukin-6 production, and chemokine production. In general, interferon-related genes showed increased accessibility, interleukin-related genes showed decreased accessibility, and chemokine-related genes varied in their direction of accessibility changes (**Figure 4A**). Upregulated DAS in innate cell types were also enriched for numerous epigenetic processes, including positive regulation of gene silencing by miRNA, negative regulation of histone H4 acetylation, histone lysine methylation, and protein K48-linked deubiquitination (**Figure 3**). Downregulated DAS were associated with downregulation of histone H3-K4 methylation, protein K63-linked deubiquitination, and peptidyl-tyrosine phosphorylation (**Figure S2**). Lysine demethylase-related genes showed increased accessibility in CD14 monocytes and decreased accessibility in CD16 monocytes and cDCs (**Figure 4B**).

**Figure 3.**
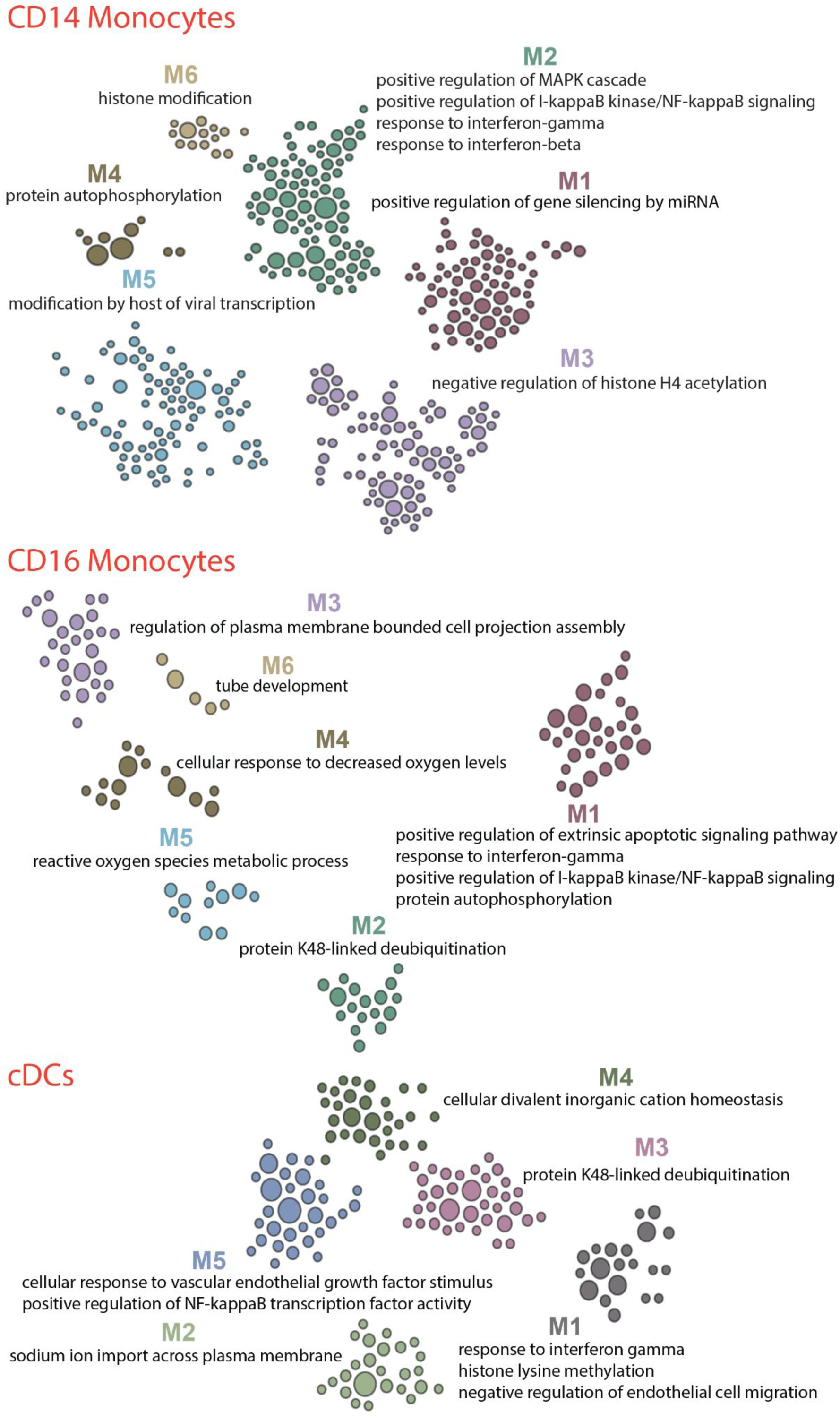
Promoter sites that are upregulated at 28 days post-challenge (versus 1 day pre-challenge) are associated with inflammatory and active epigenetic remodeling processes. (See also Figure S2.) Functional module discovery on upregulated promoter DAS from scATAC-seq data revealed modules related to inflammation and active epigenetic remodeling in classical and nonclassical monocytes as well as conventional dendritic cells (cDCs).

**Figure 4.**
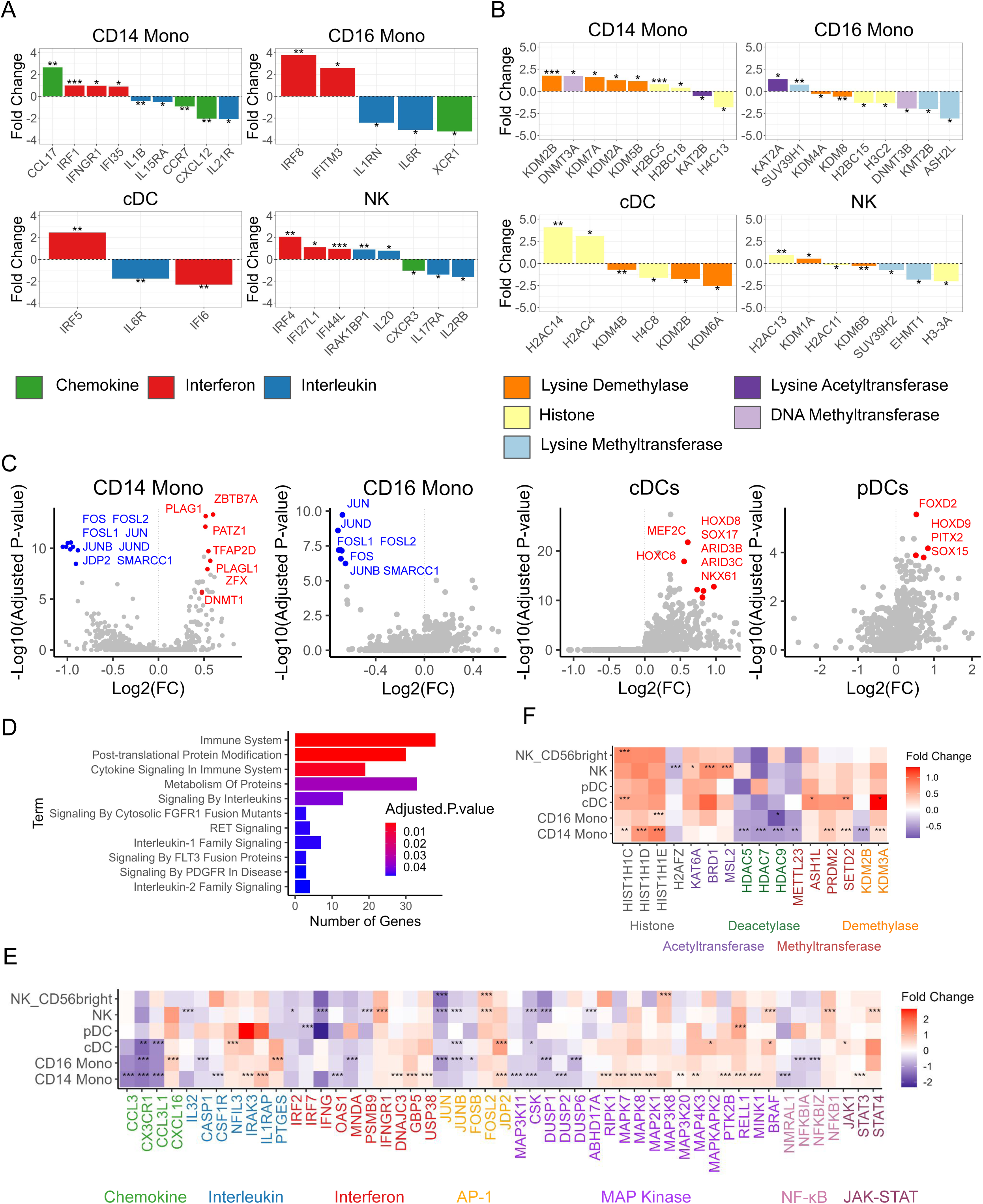
Epigenetic and transcriptional state of inflammatory and epigenetic remodeling genes. (See also Figures S3-S7 and Tables S10-11.) All analyses compared data from 28 days post-challenge to 1 day pre-challenge. (A) Statistically significant chromatin accessibility changes in promoters of cytokine-related genes in naive HVL subjects. Statistical significance is indicated with * = p-value < 0.05, ** = p-value < 0.01, *** = p-value < 0.001. (B) Statistically significant chromatin accessibility changes in promoters of epigenetic remodeling-related genes in naive HVL subjects. Statistical significance is indicated with * = p-value < 0.05, ** = p-value < 0.01, *** = p-value < 0.001. (C) Transcription factor motif enrichment analysis on differentially accessible sites from different innate immune cell types in naive HVL subjects. Downregulated motifs are indicated in blue and upregulated motifs are indicated in red. (D) Pathway enrichment analysis (Reactome) on upregulated DEG in CD14 monocytes from naive HVL subjects. (E) Heatmap of differentially expressed cytokine-related genes across different innate immune cell types in naive HVL subjects. Statistical significance is indicated with * = adjusted p-value < 0.05, ** = adjusted p-value < 0.01, *** = adjusted p-value < 0.001. (F) Heatmap of differentially expressed epigenetic remodeling-related genes across different innate immune cell types in naive HVL subjects. Statistical significance is indicated with * = adjusted p-value < 0.05, ** = adjusted p-value < 0.01, *** = adjusted p-value < 0.001.

Consonant with processes reflected by changes in chromatin accessibility, transcription factor (TF) motif enrichment analysis for upregulated and downregulated DAS in innate immune cell types revealed enrichment of numerous motifs related to inflammation and epigenetic remodeling. Both CD14 and CD16 monocytes in naive HVL subjects showed enrichment of motifs associated with activator protein 1 (AP-1) TFs (JUN, JUNB, JUND, FOS, FOSL1, and FOSL2) in downregulated DAS (**Figure 4C**). AP-1 motifs were also enriched in downregulated DAS in CD14 monocytes, cDCs, and NK cells in naive LVL subjects (**Figure S3A**) and in CD16 monocytes and NK cells in vaccinated HVL subjects (**Figure S3B**). cDCs in naive HVL subjects and pDCs in naive and vaccinated HVL subjects showed enrichment of HOX motifs, which play an important role in influencing NF-κB signaling,^21^ in upregulated DAS. In addition, CD14 monocytes in naive HVL subjects showed enrichment of key epigenetic “writer” and “reader” protein motifs in upregulated DAS, including DNMT1 and numerous zinc finger or zinc finger-like proteins (e.g., PLAG1, PATZ1, and PLAGL1).

### Transcriptional landscape reflects inflammatory and epigenetic remodeling programs

We also characterized the single-cell transcriptional programs at 28 days after infection. Reactome^22^ pathway enrichment analysis^23^ of upregulated DEG in CD14 monocytes revealed upregulation of Cytokine Signaling in Immune System, Interleukin-1 Family Signaling, and Interleukin-2 Family Signaling in naive HVL subjects (**Figure 4D**). Gene ontology analysis^23,24^ found enrichment for positive regulation of stress-activated MAPK cascade and other apoptotic processes in CD14 monocytes (**Figure S4**), overlapping with the enrichment analysis for upregulated DAS. CCL and CX3 family chemokines were downregulated, while the CXC gene CXCL16 was upregulated (**Figure 4E**). A similar trend was found in vaccinated HVL subjects, with numerous CXC family genes (CXCL2, CXCL3, and CXCL8) upregulated in CD14 monocytes (**Figure S5A**). In contrast, CXC family genes were found to be downregulated in naive LVL subjects (**Figure S5B**). We also used bulk transcriptomics (RNA-seq) to measure expression levels of chemokine genes during the acute phase of infection in naive HVL subjects (**Figure S6A** and **Table S10**) and vaccinated HVL subjects (**Figure S6B** and **Table S11**), generally finding that the direction of regulation was the opposite of what was observed at 28 days post-challenge. For instance, CXCL16 was downregulated at 8 days post-challenge in bulk data for both subject groups but upregulated in monocytes at 28 days post-challenge in the single-cell data.

Pro-inflammatory interleukins and related genes were downregulated in all subject groups at 28 days post-challenge and included IL32 and CASP1 in naive HVL subjects (**Figure 4E**), CSF1R in vaccinated HVL subjects (**Figure S5A**), and IL1B in naive LVL subjects (**Figure S5B**). Meanwhile, anti-inflammatory interleukins and related genes were upregulated, including NFIL3, IRAK3, and IL1RAP (soluble form) in naive HVL subjects and IL10, IL10RA, and IL1R2 in vaccinated HVL subjects. Like chemokines, the direction of regulation for interleukin gene expression during the acute phase of infection was largely the opposite of what was observed at 28 days post-challenge (e.g., upregulation of CASP1 and CSF1R in naive HVL subjects (**Figure S6A**) and downregulation of IL1R2 in vaccinated HVL subjects (**Figure S6B**)). Across all subject groups, most AP-1 genes (e.g., JUN, JUNB, JUND, FOS, FOSB) were found to be downregulated at 28 days post-challenge, whereas FOSL2 was generally upregulated. Importantly, the Jun dimerization protein JDP2 was found to be upregulated in naive HVL and vaccinated HVL subjects, suggesting a possible mechanism for downregulation of AP-1 activity (see Discussion). NF-κB activity was upregulated in naive HVL subjects at 28 days post-challenge, with upregulation of NFKB1 and downregulation of inhibitors NFKBIA and NFKBIZ. Finally, the transcription factor STAT3 was upregulated in CD14 monocytes in naive HVL subjects.

Antiviral activity was largely downregulated in naive HVL and vaccinated HVL subjects, with downregulation of interferon regulatory factors IRF2 and IRF7 in naive HVL subjects (**Figure 4E**) and downregulation of IRF1, IRF2, and IRF8 in vaccinated HVL subjects (**Figure S5A**). In addition, IFNG and interferon-related gene OAS1 were downregulated in both groups, while antiviral inhibitory genes USP38 and DNAJC3 were upregulated in naive HVL subjects. Notably, the interferon receptor IFNGR1 was upregulated in both groups, potentially as a means to compensate for lower interferon activity. Finally, STAT1, a key component of interferon signaling, was also found to be downregulated in vaccinated HVL subjects. Like interleukins, the direction of regulation for interferon gene expression during the acute phase of infection measured by bulk RNA-seq was largely the opposite of what was observed by single-cell analysis at 28 days post-challenge (e.g., upregulation of IRF7 and OAS1 in naive HVL subjects (**Figure S6A**) and upregulation of IFI16 and IFIH1 in vaccinated HVL subjects (**Figure S6B**)).

MAP kinase genes were generally upregulated in all four conventional MAP kinase pathways (ERK1/2, ERK5, p38, and JNK) in naive HVL and vaccinated HVL subjects at 28 days post-challenge (**Figure 4E and Figure S5A**). For instance, MAPK7 (ERK5) showed upregulation in naive HVL subjects and MAPK8 (JNK) showed upregulation in both HVL groups. Notably, MAPK14 (p38 alpha) was downregulated in vaccinated HVL subjects, but upstream gene MAP2K3 and downstream gene MAPKAPK2 were both upregulated. MAP kinase inhibitory dual specificity phosphatase (DUSP) genes were downregulated in all viral load and vaccination status subject groups.

To further examine the relationship of viral load and the post-infection immune state, we examined the correlation of the fold changes of inflammatory-related genes in the naive HVL and LVL groups. As described above, there was overlap of DEG in both HVL and LVL groups. However, comparing the fold change levels for the union of all HVL and LVL DEG provided an overall assessment of the consistency and magnitude of these post-infection gene programs following these different levels of viral infection. Overall similar response patterns were seen in most cell types, with significant correlation observed in CD14 and CD16 monocytes as well as pDCs (**Figure S7A**). The linear regressions showed a consistent slope greater than 1, indicating that while the pattern of immune-pathway regulation was shared across viral load groups, its magnitude was higher in the HVL group. A similar analysis of the naive and vaccinated HVL subjects showed correlated responses having similar magnitude across vaccination status for all cell types examined (**Figure S7B**).

Analysis of the post-challenge transcriptome also revealed differential expression of many genes associated with epigenetic remodeling in naive HVL subjects (**Figure 4F**). Expression of several histone deacetylases (HDAC5, HDAC7, and HDAC9) was downregulated in innate immune cell types, while expression of KAT6A, a histone acetyltransferase, was upregulated. Naive vaccinated subjects showed a similar trend, with downregulation of histone deacetylases HDAC2 and HDAC9 and upregulation of KAT6B. Notably, histone deacetylase gene expression was also downregulated during the acute phase of infection in naive HVL subjects (**Figure S6C**). Furthermore, histone methyltransferases (ASH1L, PRDM2, and SETD2) were generally upregulated in naive HVL subjects, and individual histone demethylases showed varying behavior (KDM2B was significantly downregulated in monocytes, while KDM3A was upregulated in most innate immune cell types). Similar to our analysis of inflammatory genes above, we examined the relationship of viral load to regulation of these epigenetic remodeling genes by comparing their fold changes in the naive HVL and LVL subjects. Regulation did not show significant correlation when comparing naive HVL and LVL transcriptome responses (**Figure S7C**). A similar analysis across vaccination status found statistically significant correlation in CD14 and CD16 monocytes as well as cDCs (**Figure S7D**).

Overall, this evidence suggests that epigenetic remodeling of the innate immune system, which is detected at 28 days post-challenge, occurs in individuals showing high viral load and reflects a complex interplay of multiple epigenetic processes.

### Gene regulatory circuits modulate expression of genes associated with inflammation and epigenetic remodeling

We next sought to identify the relationships between the post-infection regulatory events captured by each assay modality and to elucidate the multi-omic transcriptional regulatory programs they represent. The key cis-regulatory units underlying gene control involve a gene regulatory circuit (GRC) comprising the regulated gene, the cis regulatory locus, and the transcriptome factor (**Figure 5A**). We leveraged the single-cell transcriptome and chromatin data to identify GRCs that were regulated at 28 days post-challenge using the MAGICAL^25^ Bayesian framework. This framework models the regulated genes and chromatin sites using prior TF and 3D interaction data to identify GRCs at cell-type resolution. In total, we identified 443 circuits across 225 genes within innate immune cell types in naive HVL subjects (**Figure 5B** and **Table S12**), 96 circuits across 40 genes in naive LVL subjects (**Table S13**), and 802 circuits across 374 genes in vaccinated HVL subjects (**Table S14**).

**Figure 5.**
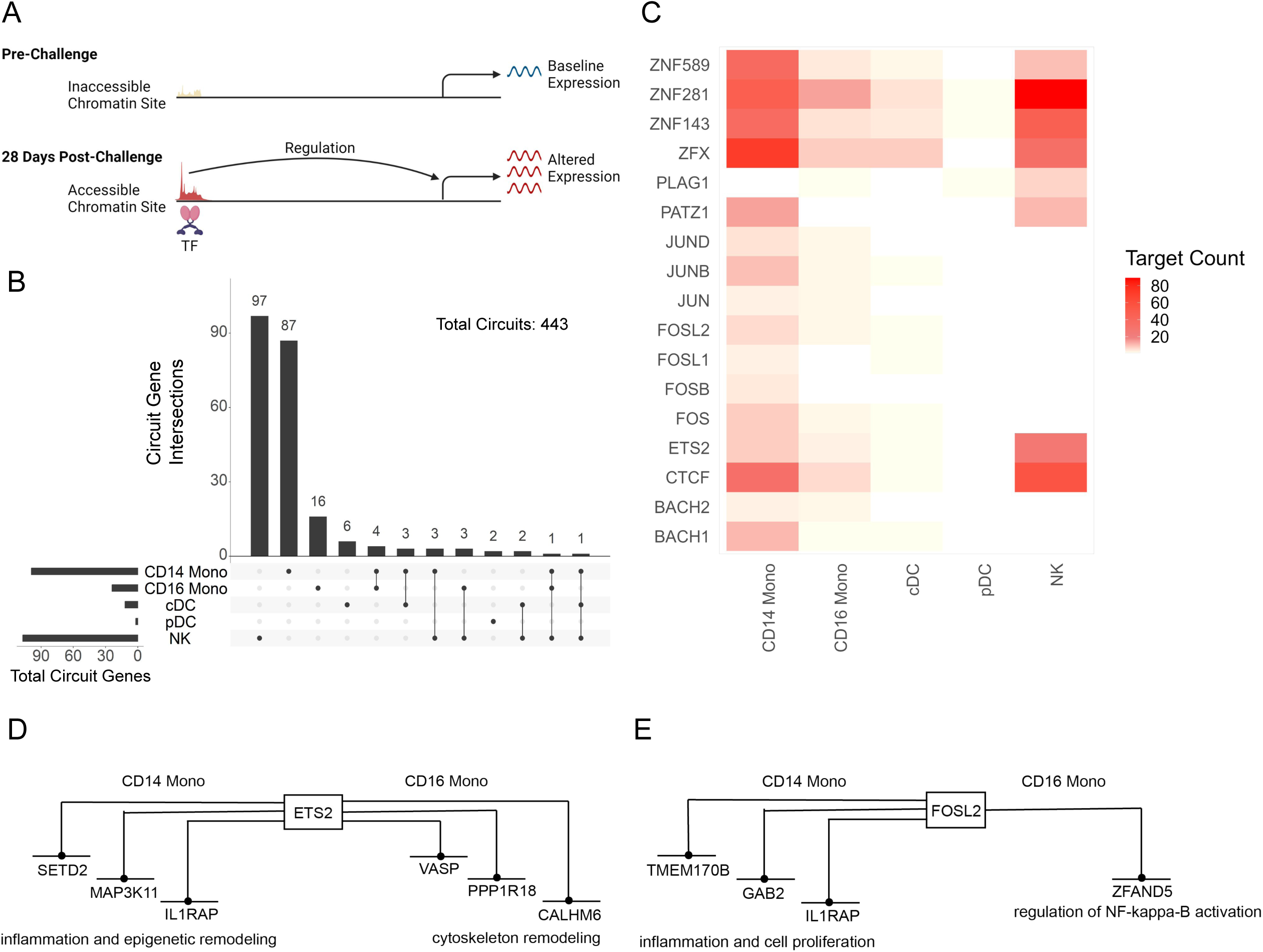
Gene regulatory circuits shape gene expression in naive HVL subjects after resolution of influenza infection. (See also Figure S8 and Tables S12-S14.) Gene regulatory circuits were found using DEG and DAS (28 days post-challenge versus 1 day pre-challenge). (A) Schematic depicting transcription factor (TF)-chromatin site-gene regulatory circuit. (B) UpSet plot showing overlap of circuit genes for innate immune cell types. (C) Heatmap of TF target counts across all gene regulatory circuits in innate immune cell types. (D) Schematic depicting gene targets of inflammatory factor ETS2. (E) Schematic depicting gene targets of AP-1 member FOSL2.

Numerous AP-1 TFs (e.g., JUN, JUNB, FOSB, FOSL1, and FOSL2) and zinc finger or zinc finger-like TFs (e.g., PATZ1, PLAG1, ZFX, and ZNF143) were found to target circuit genes in multiple innate immune cell types in naive HVL subjects (**Figure 5C**) as well as naive LVL subjects (**Figure S8A**) and vaccinated HVL subjects (**Figure S8B**). Notably, the inflammatory factor ETS2, recently implicated in directing macrophage inflammation, was found to form GRCs modulating the differential expression of several genes in CD14 and CD16 monocytes at 28 days after infection in naive HVL subjects, including the MAP kinase MAP3K11, the interleukin 1 receptor accessory protein IL1RAP, and the histone methyltransferase SETD2 (**Figure 5D**).^26^ The AP-1 member FOSL1 was also found to form GRCs in CD14 and CD16 monocytes, modulating genes related to cell proliferation and inflammation (TMEM170B, GAB2, and IL1RAP) in CD14 monocytes and the NF-κB regulator ZFAND5 in CD16 monocytes (**Figure 5E**).

To better understand the mechanistic underpinnings of inflammation and epigenetic remodeling, we examined GRCs involved in those processes. Additionally, we leveraged histone modification data from Mint-ChIP and single-nucleus methylation data to provide insight into how GRCs may interact with these other epigenetic mechanisms. Our analysis focused on CD14 monocyte GRCs in naive HVL subjects. We first evaluated the GRC modulating the upregulated MAP kinase gene MAPK7 (ERK5) (**Figure 6A**). The circuit site showed upregulated chromatin accessibility at 28 days post-challenge, and the associated TF ELF4 is well known to be involved in innate immune response and may have contributed to upregulated transcription of MAPK7. In contrast to MAPK7 and all other MAP kinase genes found to be differentially expressed in CD14 monocytes, the MAP kinase kinase kinase gene MAP3K11 showed lower expression at 28 days post-challenge versus pre-challenge (see **Figure 4E**). MAP3K11 is involved in the JNK cascade and had 4 GRCs modulating its downregulated transcription (**Figure 6B**). The inflammatory factor ETS2 showed high probability of binding with the circuit site with downregulated chromatin accessibility, indicating more limited binding of this inflammatory TF at 28 days post-challenge. At the 3 circuit sites with increased accessibility, repressive TFs like ZNF148, ZNF589, ZNF282, and E2F6 may have contributed to downregulated expression of MAP3K11. Additionally, the promoter region of MAP3K11 was hypermethylated at 28 days post-challenge and also had downregulated expression of the histone acetylation mark H3K27ac, consistent with the repression of MAP3K11. Notably, MAP3K11 showed a significant decrease in expression first detected at 8 days post-challenge versus pre-challenge (see **Figure S6A** and **Table S13**), suggesting that transcription of MAP3K11 was repressed during resolution of infection and then maintained this state at 28 days post-challenge.

**Figure 6.**
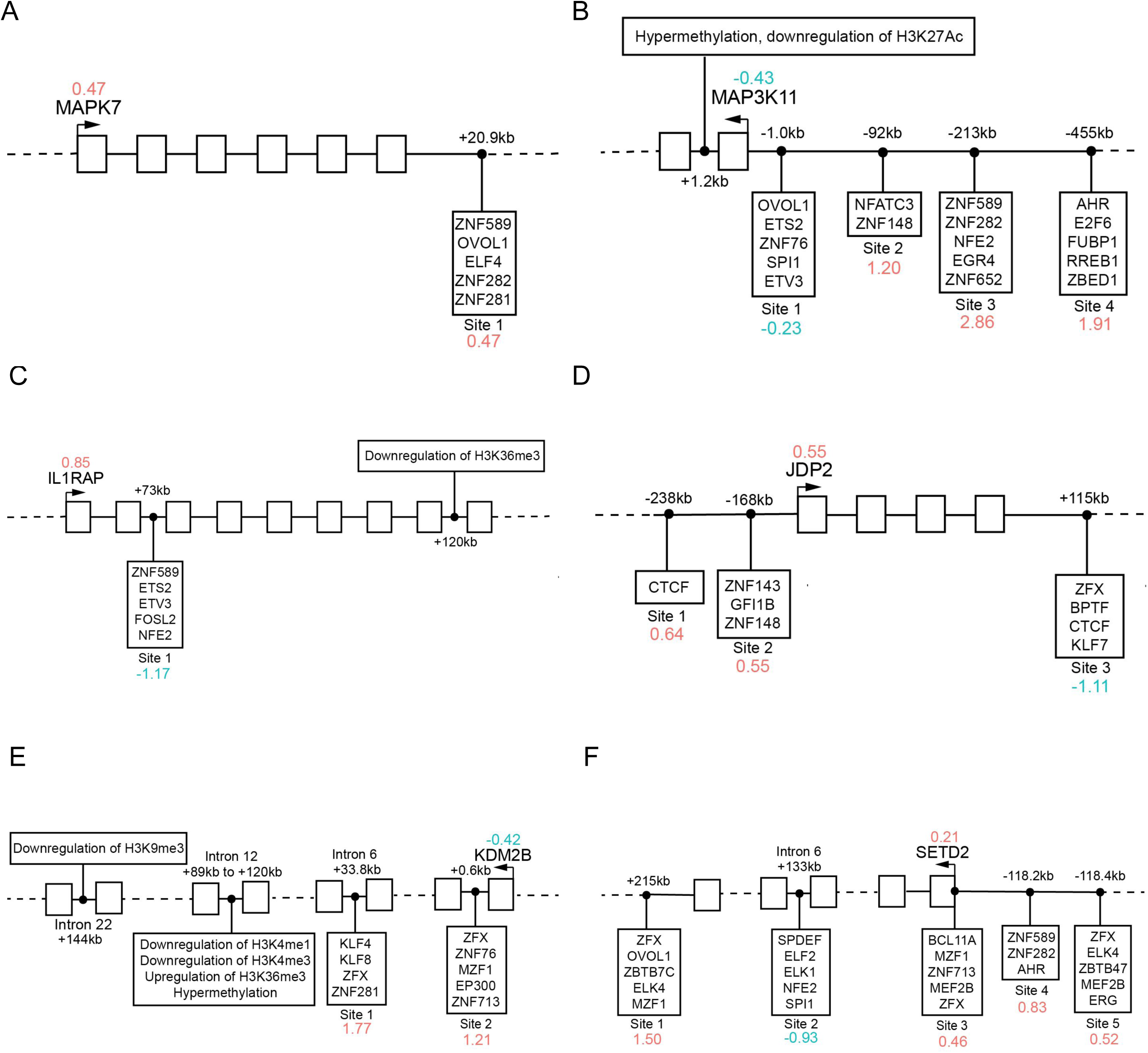
Evaluation of gene regulatory circuits in CD14 monocytes reveals important insight into the epigenetic mechanisms that control expression of inflammatory and epigenetic-remodeling genes post-influenza infection. Each schematic depicts one or more TF-chromatin site-gene regulatory circuits found using DEG and DAS (28 days post-challenge versus 1 day pre-challenge). For each chromatin site in a given circuit, all TFs that have high binding probability with the site are listed. Red values indicate upregulated fold change for a given chromatin site or gene while blue values indicate downregulated fold change. Data from other epigenetic mechanisms (DNA methylation and histone modification) are integrated into the schematic if present. (A) Schematic depicting the gene regulatory circuit that modulates MAP kinase gene MAPK7 (ERK5). (B) Schematic depicting the four gene regulatory circuits that modulate MAP kinase gene MAP3K11. In addition, the promoter region of MAP3K11 is hypermethylated and shows downregulation of the acetylation mark H3K27ac. (C) Schematic depicting the gene regulatory circuit that modulates interleukin receptor accessory protein gene IL1RAP. In addition, intron 8 of IL1RAP shows downregulation of H3K36me3. (D) Schematic depicting the three gene regulatory circuits that modulate Jun dimerization protein gene JDP2. (E) Schematic depicting the two gene regulatory circuits that modulate lysine demethylase gene KDM2B. In addition, intron 12 is hypermethylated and shows downregulation of H3K4me1, downregulation of H3K4me3, and upregulation of H3K36me3. Intron 22 shows downregulation of H3K9me3. (F) Schematic depicting the five gene regulatory circuits that modulate lysine methyltransferase SETD2.

Expression of the interleukin 1 receptor accessory protein (IL1RAP) gene, anti-inflammatory in soluble form, was elevated at 28 days post-challenge in CD14 monocytes (**Figure 6C**) and the GRC site for this gene showed decreased accessibility. While there were numerous TFs that had high probability of binding to the circuit site, ZNF589, ETV3, and NFE2 were particularly noteworthy due to their repressive roles.

Notably, the histone mark H3K36me3 was found to be downregulated in the gene body of IL1RAP. This histone mark is associated with formation of heterochromatin^19^ and could be repressing expression of IL1RAP before challenge. IL1RAP mRNA showed statistically significant downregulation at 5 and 8 days post-challenge versus pre-challenge (see **Table S13** and **Table S14**), in contrast to its upregulation at 28 days post-challenge. We also evaluated the 3 GRCs modulating the upregulated Jun dimerization protein gene JDP2 (**Figure 6D**). Transcriptional activators such as CTCF and ZNF143 may have bound to the circuit sites with increased accessibility at 28 days post-challenge, upregulating expression of JDP2. In addition, the circuit site with decreased accessibility may have limited binding of the repressive TF KLF7 at 28 days post-challenge.

There were also numerous GRCs identified regulating genes involved in epigenetic processes, such as the lysine demethylase KDM2B (**Figure 6E**) and the lysine methyltransferase SETD2 (**Figure 6F**). KDM2B expression was downregulated at 28 days post-challenge and was modulated by 2 circuit sites with upregulated chromatin accessibility as well as several other epigenetic mechanisms. Zinc-finger TFs ZNF281, ZNF76, and ZNF713 showed high probability of binding to the circuit sites, likely repressing transcription of KDM2B at 28 days post-challenge. In addition, intron 12 of KDM2B showed hypermethylation at 28 days post-challenge, downregulation of the histone mark H3K4me1 (associated with active and primed enhancers),^27^ downregulation of the histone mark H3K4me3 (associated with transcriptionally active chromatin and alternative splicing),^28^ and upregulation of the histone mark H3K36me3 (associated with formation of heterochromatin, as described above). There are a number of alternative KDM2B transcripts that have TSS located in this genomic region,^29^ and these epigenetic mechanisms may be suppressing their expression at 28 days post-challenge. In addition, downregulation of the histone mark H3K9me3 (associated with heterochromatin)^30^ was found in intron 22 of KDM2B. This repressive mark was located near another alternative KDM2B transcript TSS and may actually lead to upregulated expression of this particular transcript at 28 days post-challenge. Finally, the lysine methyltransferase SETD2 had 5 different regulatory circuits in CD14 monocytes (**Figure 6F**). The transcriptional activator ZFX showed high binding probability to 3 of the circuit sites with increased chromatin accessibility, while the ETS transcription factor ELF2 showed high binding probability to the circuit site with decreased chromatin accessibility and may indicate that transcriptional repression from this TF is removed by 28 days post-challenge.

## Discussion

In an influenza virus H3N2 challenge study, we find that mild influenza virus infection remodels the epigenomic and transcriptomic landscape of the innate immune system. These changes were detected at 28 days post-infection, weeks after clearance of the virus, and represent a post-infection modulation of inflammatory and epigenetic-remodeling programs.

The overall pattern of molecular changes observed after viral clearance were similar across viral load, particularly with respect to gene expression. For example, within nearly all innate immune cell types, while the changes in gene expression from pre-infection were generally greater in the unvaccinated subjects who had shown high viral loads, the expression changes in high and low viral load subjects were correlated (**Figure 2D**). The vaccine arm of the clinical trial failed to show an effect on viral load.^15^ Consistent with the failure of vaccination, the overall gene changes seen at 28 days after challenge in the vaccinated subjects was highly correlated with the changes observed in naive subjects (**Figure S1B**). Significant correlation in ATAC-seq data was seen in only a few cell types when comparing naive high with low viral load subjects and vaccinated with unvaccinated subjects, which most likely reflects the sparser and noisier character of this data type (**Figure S1A** and **Figure S1C**). Overall, these results suggest that the modulated epigenetic programs that persist after infection differ in magnitude but not in content in the high and low viral load subjects.

We found that innate immune system inflammatory programs were suppressed at 28 days post-infection. Inflammatory cytokine gene expression was downregulated in all subject groups (**Figure 4E**; **Figure S5A** and **Figure S5B**), while promoter regions of interleukin genes in high viral load subjects showed decreased accessibility in CD14 and CD16 monocytes as well as conventional dendritic cells (cDCs) (**Figure 4A**).

Increased activator protein-1 (AP-1) transcription factor complex activity in immune cells is associated with inflammation.^31^ In all subject groups, AP-1 genes showed decreased expression. Furthermore, the AP-1 binding site on target genes showed decreased accessibility within multiple innate immune cell types (**Figure 4C**; **Figure S3A** and **Figure S3B**). Consistent with these findings, expression of the AP-1 activity represser Jun Dimerization Protein 2 (JDP2) gene^32^ was upregulated in high viral load subjects. Furthermore, integrated single-cell gene regulatory circuit discovery found that the multivalent nuclear factor CTCF, which plays an important role in immune cell function and in the response to many viruses including influenza,^33,34^ was implicated in contributing to the regulation of JDP2 via altered accessibility of CTCF binding sites near the JDP2 gene (**Figure 6D**).

A study of epigenetic changes after influenza TIV vaccination also reported a prolonged suppression of inflammatory programs in CD14 monocytes.^13^ Unlike the decreased AP-1 accessibility we observed following influenza infection, an increased AP-1 loci accessibility in CD14 monocytes following SARS-CoV-2 infection has been reported.^35^ Various effects on innate immunity have been observed following infections or vaccinations, ranging from improved clearance of a new infection to immune dysregulation following unrelated prior immune stimuli.^3,5,36^ Overall, our results in the context of recent studies support the formulation that the epigenetic rewiring of innate immunity following an immune perturbation shows some specificity related to the particular precipitating infection or vaccination. Improving the understanding of the mechanisms underlying the alterations of innate immune memory and their functional effects should help harness this more recently recognized plasticity of innate immunity for therapeutic purposes.

Our results also show heterogeneity in the prolonged effects of immune stimuli on antiviral gene programs. We found that antiviral gene program expression was suppressed in high viral load subjects after resolution of the infection (**Figure 4E**; **Figure S5A**). These results align with the downregulated interferon response found in convalescent COVID-19 patients.^37^ In contrast with these effects in high viral load subjects, low viral load subjects had elevated antiviral gene expression at 28 days post-challenge (**Figure S5B**). Despite downregulated expression of antiviral genes, naive high viral load subjects showed upregulated accessibility in promoter regions of interferon genes in CD14 and CD16 monocytes (**Figure 4A**), and HumanBase network analysis of upregulated promoter regions in CD14 monocytes showed enrichment for “response to interferon-gamma” and “response to interferon-beta” (**Figure 2**). These results suggest that individuals who experienced high viral load may be epigenetically poised for antiviral activity, despite general suppression of inflammation.

Naive and vaccinated high viral load subjects showed overall upregulation of MAP kinase genes in CD14 monocytes at 28 days post-infection (**Figure 4E**; **Figure S5A**). For example, MAPK7 (ERK5) showed upregulation in naive high viral load subjects and MAPK8 (JNK) showed upregulation in both high viral load groups. The MAP kinase signaling cascades are involved in a wide variety of different processes, including cell proliferation and survival, cell differentiation, and apoptosis.^38^ While MAP kinase signaling is often associated with a pro-inflammatory state^39^ (e.g., through upregulation of AP-1 activity),^,40^ MAP kinase activity can also inhibit inflammation^39^ through upregulation of anti-inflammatory cytokines such as IL10 (found to be upregulated in vaccinated high viral load subjects). In addition, dual specificity phosphatase (DUSP) genes were downregulated in all subject groups across multiple innate immune cell types. DUSP enzymes inactivate MAP kinase genes through dephosphorylation of threonine and tyrosine residues in their activation loops.^39^ DUSP1-knockout mice showed increased production of pro-inflammatory cytokines (e.g., IL1B and IL6) relative to wild type mice in macrophages after infection with bacteria Chlamydia pneumoniae.^41^ Thus, despite inflammatory cytokine and AP-1 activity being suppressed as described above, upregulated MAP kinase activity and downregulated DUSP activity may suggest that the innate immune system is primed to respond strongly to secondary immune challenges after resolution of influenza infection.

We also found evidence of epigenetic-remodeling immune programs at 28 days post-infection. Notably, histone deacetylation genes were downregulated in CD14 monocytes in both naive and vaccinated high viral load subjects (**Figure 4F**), while histone acetyltransferase KAT6B was upregulated in vaccinated high viral load subjects (**Table S8**). Convalescent COVID patients from a previous study conducted by Cheong et al. showed a similar trend, with upregulation of KAT6A and downregulation of HDAC8 and HDAC9 in monocytes 2 to 4 months after onset of disease.^35^ In contrast, subjects vaccinated with TIV showed opposite behavior (downregulation of histone acetyltransferase genes and upregulation of histone deacetylase genes) at 7 days after vaccination. Using bulk transcriptomics (RNA-seq) data, we found downregulation of HDAC5 and HDAC7 at 5 days and 8 days post-infection in naive high viral load subjects (**Figure S6C**).^13^ This suggests that histone acetylation activity after influenza infection may more closely resemble the immune state after SARS-CoV–2 infection versus TIV vaccination. We also found that CD14 monocytes showed increased chromatin accessibility at lysine demethylases promoter sites, while CD16 monocytes and cDCs showed decreased chromatin accessibility (**Figure 4B**). This may reflect the specific roles of these innate immune cell types: CD14 monocytes may be poised to produce lysine demethylases and activate genes when secondary infection occurs, whereas CD16 monocytes and cDCs may be more stable.

In conclusion, our results demonstrate that mild influenza virus infection causes persistent epigenetic and transcriptional changes to the innate immune system that are present 28 days post-infection. These changes comprise a complex alteration in the expression levels and chromatin state of inflammatory and antiviral gene programs.

These findings support the emerging view of innate immune system plasticity and that the patterns of innate immune system adaptations are dependent on the specific immune perturbation.

## Limitations of the study

The study being a controlled challenge provided unique benefits, but the results observed may differ from that seen in natural infection. The design of our study does not allow the functional effects of the molecular adaptations of innate immunity identified to be experimentally determined.

## Acknowledgments

We thank the Single-cell and Spatial Technologies team at the Center for Advanced Genomics Technology, Department of Genetics and Genomic Sciences, the Icahn School of Medicine at Mount Sinai for providing expertise on experimental and computational matters. We are grateful for funding from the NIH NIGMS grant no. R01GM071966 (O.G.T.), NHGRI grant no. R01HG005998 (O.G.T.), NIH NHGRI training grant T32HG003284 (O.G.T.), Simons Foundation grant no. 395506 (O.G.T.), NIH NIDDK grant no. R01DK46943 (S.C.S.), and Defense Advanced Research Projects Agency contract no. N6600119C4022 (S.C.S.). BioRender.com was used to create Figures 1A, 1B, and 5A.

## Author contributions

M.T.M., T.W.B., S.H.K., J.R.E., C.W.W., W.J.G., X.C., I.R., E.Z., T.G.E., S.C.S., and O.G.T. conceived the study and supervised the research. W.T., W.S.C., P.A., A.R., and D.C. conducted computational analyses. S.V. collected and processed the single-cell RNA-seq and ATAC-seq data. P.A., R.M., and C.W.W. processed the Mint-ChIP data. W.W., B.W., M.H., J.R.E., and W.J.G. processed the snMethylation data. W.T., S.V., W.S.C., A.C., X.C., I.R., E.Z., S.C.S., and O.G.T. wrote the original draft of the manuscript. All authors proofread the submitted version.

## Methods

### Lead contact

Further information and requests for resources should be directed to and will be fulfilled by the lead contact, Stuart C. Sealfon.

### Materials availability

This study did not generate new materials.

### Data availability

Samples were collected in collaboration with the Biomedical Advanced Research and Development Authority (BARDA) office and raw data will be released through the Defense Threat Reduction Agency (DTRA) in the near future (as the associated Epigenetic CHaracterization and Observation (ECHO) effort from the Defense Advanced Research Projects Agency (DARPA) concluded in August 2024). We do not yet have accessions for access to the raw data but will make them available to reviewers as soon as possible. Processed data are available via the supplemental tables.

### Code availability

Analysis code can be found at the following GitHub repository: https://github.com/williamthistle/Influenza

### Study outline

The samples used in this study were generated from a phase 2, single center, randomized, double blind clinical trial (https://clinicaltrials.gov/study/NCT03883113). Healthy, non-smoking adult volunteers between the ages of 18 and 55 who had not received the 2018-19 influenza vaccine or any other live vaccine in the 4 weeks prior to enrollment, had microneutralization titers <20 against the influenza virus A/Belgium/4217/2015 (H3N2) strain, and were willing to remain in isolation for the duration of the study were included. 819 subjects were screened, 145 subjects were enrolled and vaccinated (out-patient phase, randomized 87:58 vaccine:placebo), and 117 subjects were subsequently challenged at least 6 weeks later and completed the study (in-patient phase, at an SGS quarantine facility in Antwerp, Belgium).

Vaccinated individuals received 0.5mL (1.5 x 10^8 pfu) of the MVA-NP+M1 vaccine (derived from H3N2/Panama/2007; Emergent Biosciences, Rockville, MD, USA) intramuscularly while placebo group individuals received 0.5mL of saline (0.9% NaCl). The challenge agent, influenza virus H3N2 A/Belgium/4217/2015 (Meridian Life Science, Memphis, TN, USA) was administered to all subjects (0.25 ml in each nostril, 1.0×10^6 TCID50/ml) via a nasal spray (Teleflex MAD301 device, Morrisville, NC, USA). The primary outcome for the trial was the reduction of nasopharyngeal viral shedding.

Nasal swabs were taken from subjects twice a day (at least 8 hours apart) for 10 consecutive days, including the day of challenge. Viral load (measured by qRT-PCR) was calculated from cumulative area under the curve (AUC) plotted using log viral particle number/ml for each time point against time. The secondary outcome included comparison of incidence of laboratory confirmed (culture/qRT-PCR) influenza-like-illness (ILI), the AUC of total symptom scores, days of fever, mucus weight, correlation of T cell responses (ELISPOT) to viral load AUC, and the attack rate (% of challenged subjects with at least 2 qRT-PCR positive swabs), between the MVA-NP+M1 (“vaccinated”) and placebo (“naive”) groups. At the end of the trial period, the vaccine did not induce significant reduction in viral load or symptoms.^15^ Both primary and secondary outcome measures were utilized in sample selection for this study.

### Experimental protocols

Venous blood samples were collected into Sodium Heparin CPT tubes (BD) for PBMC generation (at Viroclinics, cryopreserved using CTL Cryo-ABC kit, CTL-Europe, Bonn, Germany) and PAXgene® RNA tubes (for transcriptional analysis). Nasal swabs were collected into viral transport medium, frozen, and shipped to Viroclinics (Rotterdam, Netherlands). A positive qRT-PCR was verified by quantitative culture TCID50 using MDCK cells. Cryopreserved PBMCs were thawed and used for measuring T cell responses against NP and M1 peptides with two assays- 1) an IFNγ and Granzyme B ELISPOT assay (CTL-Europe, Bonn, Germany), and 2) intracellular cytokine staining for IFNγ, TNFα and IL-2 using flow cytometry (Caprion, Ghent, Belgium).

### Sample selection

The primary goal of the study was to assess epigenetic and transcriptional changes following mild influenza infection. Study subjects who received the placebo and were later challenged with live virus were used to model immune responses as “naive” individuals who experienced a mild influenza virus infection, whereas “vaccinated” individuals provided a well-matched population for comparison. A total of 45 (19 HVL, 10 MVL, 16 LVL) placebo subjects and 69 MVA-NP+M1 vaccinated subjects (22 HVL, 25 MVL, 22 LVL) were chosen for this study. All 114 virus-challenged subjects were first ranked by viral load AUC to stratify subjects into those who developed high viral load (AUC ≥ 959, HVL), moderate viral load (AUC > 458 and AUC < 959, MVL) or low viral load (AUC ≤ 458, LVL). PBMCs at 1 day pre-challenge and 28 days post challenge from 23 naive subjects (13 HVL, 10 LVL), and 14 vaccinated subjects (7 with the highest and 7 with the lowest magnitude of T cell responses to NP and M1 peptides at Day 28 following vaccination) were selected for single-cell RNA-Seq and single-cell ATAC-Seq assays. Only a portion of samples passed QC and were used in analysis (exact numbers detailed in **Table S1** and **Table S2**). PBMCs from the 13 naive high viral load individuals were also chosen for single-nucleus DNA methylation analysis (performed at the Salk Institute)^16^, and PBMCs from 11 naive high viral load individuals were chosen for multiplexed, indexed T7 ChIP-seq (Mint-ChIP) analysis to measure histone levels for 6 different histone features (performed at Duke University, see details below). Finally, PAXgene® samples from all 114 challenged subjects (45 naive + 69 vaccinated) at 6 time points (2 days and 1 day pre-challenge; and 2, 5, 8 and 28 days post-challenge, a total of 289 samples) were subjected to bulk transcriptomic (RNA-seq) analysis.

### Bulk RNA-seq data processing and analysis

Reads from all samples were processed using the MoTrPAC RNA-Seq pipeline,^42^ resulting in a transcript-by-sample unnormalized count matrix. Data were first separated into each experimental period (vaccination phase and influenza challenge phase) and then separated by viral load (low, moderate, and high) for further downstream analysis.

Within each experimental period and viral load group, differential expression analysis was performed using the R package DESeq2.^43^ In all differential expression analyses, we integrated surrogate variables into our statistical models using the R package SVA^44^ to help control for false positives due to cell type proportion changes and technical variation between samples. Wald tests were performed to detect differentially expressed genes (DEGs) between pre-vaccination samples and 2, 8, and 28 days post-vaccination samples. In addition, Wald tests were performed to detect DEGs between pre-exposure to influenza samples and 2, 5, 8, and 28 days post-exposure to influenza samples. In all cases, adjusted p-value < 0.05 (alpha = 0.05) was used as a significance threshold, and different log2 fold change thresholds (lfcThreshold = 0.1, 0.2, 0.3, 0.585, 1, or 2) were used to capture DEGs with varying levels of stringency.

### scRNA-seq data processing and analysis

Briefly, cryopreserved PBMCs were thawed with warm RPMI (Gibco, Grand Island, United States) + 10% FBS (Thermo Fisher, Waltham, United States), and washed with PBS (Gibco, Grand Island, United States) + 0.4% BSA (Sigma Aldrich, St. Louis, United States). Reads from all samples were aligned to human reference genome GRCh38 (hg38) using 10x Genomics Cell Ranger v6.1.0. Filtered feature-barcode matrices were then processed using the Single-cell Pipeline for End to End Data Integration (SPEEDI) pipeline.^45^ SPEEDI performed QC filtering with dynamically chosen thresholds, data-driven batch identification, data integration, and cell-type labeling. We used a PBMC reference object generated by the Satija lab for cell type annotation.^46^ Data were then separated by vaccination status (mock-vaccination and MVA-NP+M1 vaccination) and viral load (low and high) for further downstream analysis.

Within each cell type, differentially expressed genes (DEG) were found for our chosen contrast (28 days post-exposure to influenza vs pre-exposure to influenza) using a two-step process. First, differential expression analysis was performed on the single-cell level using Seurat v5.^47^ We used the FindMarkers function, with thresholds |log2FC| ≥ 0.1, adjusted p-value < 0.05, and min.pct threshold of 0.1 (genes must be expressed in at least 10% of cells in the pre- or post-exposure group). Importantly, because each subject provided both pre- and post-exposure samples, we used a logistic regression approach (test.use = “LR” in FindMarkers) and included the subject ID as a covariate to increase power. Next, to correct for the high false positive rate usually found in single-cell DEG discovery, we aggregated single-cell gene expression data into cell-type level pseudobulk and performed differential expression analysis using DESeq2. As above, we included subject ID in our model to increase power. Genes that had p-value < 0.05 (Wald test) and were also found to be statistically significant in the single-cell analysis were selected as the final set of differentially expressed genes.

For evaluation of biological processes associated with DEGs within each cell type, we used Enrichr^23^ to perform both Gene Ontology (GO)^24^ enrichment analysis and pathway enrichment analysis with the Reactome Knowledgebase.^22^ We split DEGs according to the direction of fold change (positive or negative) for analysis.

### scATAC-seq data processing and analysis

Cryopreserved PBMCs were processed as described above for scRNA-seq. Reads from all samples were aligned to human reference genome GRCh38 (hg38) using 10x Genomics Cell Ranger ATAC v2.0.0. Filtered ATAC fragment files were then processed using SPEEDI and the R package ArchR.^48^ SPEEDI was used to read in the data and remove low quality cells, including doublets. Next, the ArchR function addIterativeLSI was used to perform dimensionality reduction on the data. Cell types were annotated using ArchR’s addGeneIntegrationMatrix function, using the same PBMC reference as the scRNA-seq analysis. Peaks were called with MACS2 via ArchR’s addReproduciblePeakSet function to create a union reproducible peak set. After processing, data were separated by vaccination status (mock-vaccination and MVA-NP+M1 vaccination) and viral load (low and high) for further downstream analysis.

Next, for differentially accessibility analysis, we converted the ArchRProject object into a Signac^49^ SeuratObject using the ArchRtoSignac R package.^50^ Within each cell type, differentially accessible sites (DASs) were found for our chosen contrast (28 days post-exposure to influenza vs pre-exposure to influenza) using a two-step process. First, differential accessibility analysis was performed on the single-cell level using Seurat’s FindMarkers function^4^, with thresholds |log2FC| ≥ 0.1, p-value < 0.05, and min.pct threshold of 0.01 (sites must be expressed in at least 1% of cells in the pre- or post-exposure group). Importantly, because each subject provided both pre- and post-exposure samples, we used a logistic regression approach (test.use = “LR” in FindMarkers) and included the subject ID as a covariate to increase power. We also included nPeaks as a covariate to control for the total number of fragments in a given cell. To correct for the high false positive rate usually found in single-cell DAS discovery, we aggregated single-cell data into cell-type level pseudobulk and performed differential accessibility analysis using two different linear model approaches. We first applied a simple linear model (using the lm function in R) and then applied a robust linear model (using the rlm function from the MASS^51^ R package and the f.robftest function from the sfsmisc^52^ R package) to see whether accessibility was different between the pre-exposure and post-exposure groups for a given site. Sites that had p-value < 0.05 or robust p-value < 0.05 and were also found to be statistically significant in the single-cell analysis were selected as the final set of differentially accessible sites. Sites that passed the single-cell accessibility test but not the cell-type level pseudobulk test were used for TF analysis and for generation of MAGICAL circuits. The R package ChIPseeker^53^ was used to annotate sites (using the human genome annotation packages org.Hs.eg.db^54^ and TxDb.Hsapiens.UCSC.hg38.knownGene).^55^ For promoter-specific analyses, we subset annotated sites to the “Promoter (<=1kb)”, “Promoter (1-2kb)”, and “Promoter (2-3kb)” categories.

We used the “cisbp” motif set from the chromVAR package for motif annotation.^56^ Motifs were added to the Signac object using the AddMotifs function from the Signac R package. Next, for each cell type, sites were split into upregulated and downregulated groups. For a given group of sites, a background set of 40000 sites from the same cell type were chosen based on GC content. Finally, we used the FindMotifs function from the Signac R package to find significant motifs. We used an adjusted p value of 0.05 as our significance threshold. A variety of fold change thresholds (0.1, 0.3, 0.585, 1, and 2) were tested for each cell type.

For evaluation of biological processes associated with DAS within each cell type, we used HumanBase to perform functional module discovery by mapping each DAS to its closest gene. In brief, functional module discovery uses a shared k-nearest-neighbors and Louvain community-finding algorithmic approach to cluster genes together based on a tissue-specific background interaction network.^20^ We split DAS according to the direction of fold change (positive or negative) for analysis. In all cases, the “blood” interaction network was used.

### Gene regulatory circuit (GRC) data processing and analysis

For gene regulatory circuit analysis, MAGICAL^25^ was used. DEG from the scRNA-seq analysis (28 days post-challenge versus 1 day pre-challenge) were used as candidate genes, and DAS from the scATAC-seq analysis (28 days post-challenge versus 1 day pre-challenge) were used as candidate sites. We used the topologically associating domain (TAD) boundaries file and the hg38 RefSeq file included with the MAGICAL R script. Default settings were used for processing. Circuit gene overlap counts were visualized using the R package UpSetR.^57^

### Mint-ChIP data processing and analysis

The experimental protocol used for Mint-ChIP sample generation can be found in Schuetter et al.^58^ FASTQ files were processed through the ENCODE pipeline to produce BAM files.^59^ Profiled marks included H3K4me1, H3K4me3, H3K9me3, H3K27ac, H3K27me3, and H3K36me3. For each mark, differentially enriched peaks were found for our chosen contrast (28 days post-exposure to influenza vs pre-exposure to influenza) using DiffBind^60^ and DESeq2. Because each subject provided both pre- and post-exposure samples, we included subject ID in our DESeq2 model to increase power. The p-value < 0.05 was used as a significance threshold (Wald test), and different log2 fold change thresholds (lfcThreshold = 0, 0.1, 0.2, 0.3, 0.585, 1, or 2) were used to capture differential histone marks with varying levels of stringency. The R package ChIPseeker was used to annotate peaks (using the human genome annotation packages org.Hs.eg.db and TxDb.Hsapiens.UCSC.hg19.knownGene).^61^ For integrative analyses with data from other assays, the UCSC LiftOver tool was used to convert hg19 coordinates to corresponding hg38 coordinates.^62^

## References

1. de Laval B, Maurizio J, Kandalla PK, Brisou G, Simonnet L, Huber C, Gimenez G, Matcovitch-Natan O, Reinhardt S, David E, Mildner A. C/EBPβ-dependent epigenetic memory induces trained immunity in hematopoietic stem cells. Cell stem cell. 2020 May 7;26(5):657–74. 10.1016/j.stem.2020.03.014

2. Dagenais, Amy, Carlos Villalba-Guerrero, and Martin Olivier. “Trained immunity: A “new” weapon in the fight against infectious diseases.” Frontiers in Immunology 14 (2023): 1147476. 10.3389/fimmu.2023.1147476

3. Lefkowitz, Rivka Bella, et al. “Epigenetic Control of Innate Immunity: Consequences of Acute Respiratory Virus Infection.” Viruses 16.2 (2024): 197. 10.3390/v16020197

4. Schulte-Schrepping, Jonas, et al. “Severe COVID-19 is marked by a dysregulated myeloid cell compartment.” Cell 182.6 (2020): 1419–1440. 10.1016/j.cell.2020.08.001

5. Mao, Weiguang, et al. “A methylation clock model of mild SARS-CoV-2 infection provides insight into immune dysregulation.” Molecular Systems Biology 19.5 (2023): e11361. 10.15252/msb.202211361

6. Pischedda, Sara, et al. “Role and diagnostic performance of host epigenome in respiratory morbidity after RSV infection: the EPIRESVi study.” Frontiers in immunology 13 (2022): 875691. 10.3389/fimmu.2022.875691

7. Aegerter, Helena, et al. “Influenza-induced monocyte-derived alveolar macrophages confer prolonged antibacterial protection.” Nature immunology 21.2 (2020): 145–157. 10.1038/s41590-019-0568-x

8. Sparks, Rachel, et al. “Influenza vaccination reveals sex dimorphic imprints of prior mild COVID-19.” Nature 614.7949 (2023): 752–761. 10.1038/s41586-022-05670-5

9. Morens, David M., Jeffery K. Taubenberger, and Anthony S. Fauci. “Predominant role of bacterial pneumonia as a cause of death in pandemic influenza: implications for pandemic influenza preparedness.” The Journal of infectious diseases 198.7 (2008): 962–970. 10.1086/591708

10. Jochems, Simon P., et al. “Inflammation induced by influenza virus impairs human innate immune control of pneumococcus.” Nature immunology 19.12 (2018): 1299–1308. 10.1038/s41590-018-0231-y

11. Zhao, Zhiyan, et al. “Host DNA Demethylation Induced by DNMT1 Inhibition Up-Regulates Antiviral OASL Protein during Influenza a Virus Infection.” Viruses 15.8 (2023): 1646. 10.3390/v15081646

12. Nagesh, Prashanth Thevkar, and Matloob Husain. “Influenza A virus dysregulates host histone deacetylase 1 that inhibits viral infection in lung epithelial cells.” Journal of virology 90.9 (2016): 4614–4625. 10.1128/jvi.00126-16

13. Wimmers, Florian, et al. “The single-cell epigenomic and transcriptional landscape of immunity to influenza vaccination.” Cell 184.15 (2021): 3915–3935. 10.1016/j.cell.2021.05.039

14. Evans, Thomas G., et al. “Efficacy and safety of a universal influenza A vaccine (MVA-NP+ M1) in adults when given after seasonal quadrivalent influenza vaccine immunisation (FLU009): a phase 2b, randomised, double-blind trial.” The Lancet Infectious Diseases 22.6 (2022): 857–866. 10.1016/S1473-3099(21)00702-7

15. Evans, Thomas G., et al. “Assessment of CD8+ T-cell mediated immunity in an influenza A (H3N2) human challenge model in Belgium: a single centre, randomised, double-blind phase 2 study.” The Lancet Microbe (2024). 10.1016/S2666-5247(24)00024-7

16. Wang, Wenliang, et al. “Human Immune Cell Epigenomic Signatures in Response to Infectious Diseases and Chemical Exposures.” bioRxiv (2023). 10.1101/2023.06.29.546792

17. van Galen, Peter, et al. “A multiplexed system for quantitative comparisons of chromatin landscapes.” Molecular cell 61.1 (2016): 170–180. 10.1016/j.molcel.2015.11.003

18. Zhao, Weiye, et al. “Investigating crosstalk between H3K27 acetylation and H3K4 trimethylation in CRISPR/dCas-based epigenome editing and gene activation.” Scientific reports 11.1 (2021): 15912. 10.1038/s41598-021-95398-5

19. Hahn, Maria A., et al. “Relationship between gene body DNA methylation and intragenic H3K9me3 and H3K36me3 chromatin marks.” PloS one 6.4 (2011): e18844. 10.1371/journal.pone.0018844

20. Krishnan, Arjun, et al. “Genome-wide prediction and functional characterization of the genetic basis of autism spectrum disorder.” Nature neuroscience 19.11 (2016): 1454–1462. 10.1038/nn.4353

21. Pai, Priya, and Saraswati Sukumar. “HOX genes and the NF-κB pathway: a convergence of developmental biology, inflammation and cancer biology.” Biochimica et Biophysica Acta (BBA)-Reviews on Cancer 1874.2 (2020): 188450. 10.1016/j.bbcan.2020.188450

22. Milacic, Marija, et al. “The reactome pathway knowledgebase 2024.” Nucleic acids research 52.D1 (2024): D672–D678. 10.1093/nar/gkad1025

23. Chen, Edward Y., et al. “Enrichr: interactive and collaborative HTML5 gene list enrichment analysis tool.” BMC bioinformatics 14 (2013): 1–14. 10.1186/1471-2105-14-128

24. Ashburner, Michael, et al. “Gene ontology: tool for the unification of biology.” Nature genetics 25.1 (2000): 25–29. 10.1038/75556

25. Chen, Xi, et al. “Mapping disease regulatory circuits at cell-type resolution from single-cell multiomics data.” Nature computational science 3.7 (2023): 644–657. 10.1038/s43588-023-00476-5

26. Stankey, C. T., et al. “A disease-associated gene desert directs macrophage inflammation through ETS2.” Nature (2024): 1–10. 10.1038/s41586-024-07501-1

27. Rada-Iglesias, Alvaro. “Is H3K4me1 at enhancers correlative or causative?.” Nature genetics 50.1 (2018): 4–5. 10.1038/s41588-017-0018-3

28. Beacon, Tasnim H., et al. “The dynamic broad epigenetic (H3K4me3, H3K27ac) domain as a mark of essential genes.” Clinical epigenetics 13 (2021): 1–17. 10.1186/s13148-021-01126-1

29. Martin, Fergal J., et al. “Ensembl 2023.” Nucleic acids research 51.D1 (2023): D933–D941. 10.1093/nar/gkac958

30. Nicetto, Dario, and Kenneth S. Zaret. “Role of H3K9me3 heterochromatin in cell identity establishment and maintenance.” Current opinion in genetics & development 55 (2019): 1–10. 10.1016/j.gde.2019.04.013

31. Bhosale, Pritam Bhagwan, et al. “Structural and functional properties of activator protein-1 in cancer and inflammation.” Evidence-Based Complementary and Alternative Medicine 2022.1 (2022): 9797929. 10.1155/2022/9797929

32. Aronheim, Ami, et al. “Isolation of an AP-1 repressor by a novel method for detecting protein-protein interactions.” Molecular and cellular biology 17.6 (1997): 3094–3102. 10.1128/MCB.17.6.3094

33. Nikolic, Tatjana, et al. “The DNA-binding factor Ctcf critically controls gene expression in macrophages.” Cellular & molecular immunology 11.1 (2014): 58–70. 10.1038/cmi.2013.41

34. Li, Yuchang, et al. “The critical role of human transcriptional repressor CTCF mRNA up-regulation in the induction of anti-HIV-1 responses in CD4+ T cells.” Immunology letters 117.1 (2008): 35–44. 10.1016/j.imlet.2007.11.017

35. Cheong, Jin-Gyu, et al. “Epigenetic memory of coronavirus infection in innate immune cells and their progenitors.” Cell 186.18 (2023): 3882–3902. 10.1016/j.cell.2023.07.019

36. Netea, Mihai G., et al. “Defining trained immunity and its role in health and disease.” Nature Reviews Immunology 20.6 (2020): 375–388. 10.1038/s41577-020-0285-6

37. Liu, Zhaoli, et al. “Multi-omics integration reveals only minor long-term molecular and functional sequelae in immune cells of individuals recovered from COVID-19.” Frontiers in Immunology 13 (2022): 838132. 10.3389/fimmu.2022.838132

38. Zhang, Wei, and Hui Tu Liu. “MAPK signal pathways in the regulation of cell proliferation in mammalian cells.” Cell research 12.1 (2002): 9–18. 10.1038/sj.cr.7290105

39. Arthur, J. Simon C., and Steven C. Ley. “Mitogen-activated protein kinases in innate immunity.” Nature Reviews Immunology 13.9 (2013): 679–692. 10.1038/nri3495

40. Whitmarsh, A. J., and R. J. Davis. “Transcription factor AP-1 regulation by mitogen-activated protein kinase signal transduction pathways.” Journal of molecular medicine 74 (1996): 589–607. 10.1007/s001090050063

41. Rodriguez, Nuria, et al. “Increased inflammation and impaired resistance to Chlamydophila pneumoniae infection in Dusp1-/-mice: critical role of IL-6.” Journal of leukocyte biology 88.3 (2010): 579–587. 10.1189/jlb.0210083

42. Sanford, James A., et al. “Molecular transducers of physical activity consortium (MoTrPAC): mapping the dynamic responses to exercise.” Cell 181.7 (2020): 1464–1474. 10.1016/j.cell.2020.06.004

43. Love, Michael I., Wolfgang Huber, and Simon Anders. “Moderated estimation of fold change and dispersion for RNA-seq data with DESeq2.” Genome biology 15 (2014): 1–21. 10.1186/s13059-014-0550-8

44. Leek, Jeffrey T., et al. “The sva package for removing batch effects and other unwanted variation in high-throughput experiments.” Bioinformatics 28.6 (2012): 882–883. 10.1093/bioinformatics/bts034

45. Wang, Yuan, et al. “Automated single-cell omics end-to-end framework with data-driven batch inference.” bioRxiv (2023). 10.1101/2023.11.01.564815

46. Hao, Yuhan, et al. “Integrated analysis of multimodal single-cell data.” Cell 184.13 (2021): 3573–3587. 10.1016/j.cell.2021.04.048

47. Hao, Yuhan, et al. “Dictionary learning for integrative, multimodal and scalable single-cell analysis.” Nature biotechnology 42.2 (2024): 293–304. 10.1038/s41587-023-01767-y

48. Granja, Jeffrey M., et al. “ArchR is a scalable software package for integrative single-cell chromatin accessibility analysis.” Nature genetics 53.3 (2021): 403–411. 10.1038/s41588-021-00790-6

49. Stuart, Tim, et al. “Single-cell chromatin state analysis with Signac.” Nature methods 18.11 (2021): 1333–1341. 10.1038/s41592-021-01282-5

50. Shi, Zechuan, et al. “Protocol for single-nucleus ATAC sequencing and bioinformatic analysis in frozen human brain tissue.” STAR protocols 3.3 (2022): 101491. 10.1016/j.xpro.2022.101491

51. 51. Venables, W. N., and B. D. Ripley. “Modern Applied Statistics with S, Springer, New York: ISBN 0-387-95457-0.” (2002). 10.1007/978-0-387-21706-2

52. Maechler, Martin, et al. “Package ‘sfsmisc’.” (2024). 10.32614/CRAN.package.sfsmisc

53. Yu, Guangchuang, Li-Gen Wang, and Qing-Yu He. “ChIPseeker: an R/Bioconductor package for ChIP peak annotation, comparison and visualization.” Bioinformatics 31.14 (2015): 2382–2383. 10.1093/bioinformatics/btv145

54. Carlson M (2019). org.Hs.eg.db: Genome wide annotation for Human. R package version 3.8.2. doi:10.18129/B9.bioc.org.Hs.eg.db

55. Team BC, Maintainer BP (2019). TxDb.Hsapiens.UCSC.hg38.knownGene: Annotation package for TxDb object(s). R package version 3.4.6. doi:10.18129/B9.bioc.TxDb.Hsapiens.UCSC.hg38.knownGene

56. Schep, Alicia N., et al. “chromVAR: inferring transcription-factor-associated accessibility from single-cell epigenomic data.” Nature methods 14.10 (2017): 975–978. 10.1038/nmeth.4401

57. Conway, Jake R., Alexander Lex, and Nils Gehlenborg. “UpSetR: an R package for the visualization of intersecting sets and their properties.” Bioinformatics 33.18 (2017): 2938–2940. 10.1093/bioinformatics/btx364

58. Schuetter, Jared, et al. “Integrated epigenomic exposure signature discovery.” Epigenomics (2024): 1–17. 10.1080/17501911.2024.2375187

59. Hitz, Benjamin C., et al. “The ENCODE uniform analysis pipelines.” Research Square (2023). 10.21203/rs.3.rs-3111932/v1

60. Ross-Innes, Caryn S., et al. “Differential oestrogen receptor binding is associated with clinical outcome in breast cancer.” Nature 481.7381 (2012): 389–393. 10.1038/nature10730

61. Carlson M, Maintainer BP (2015). TxDb.Hsapiens.UCSC.hg19.knownGene: Annotation package for TxDb object(s). R package version 3.2.2. doi:10.18129/B9.bioc.TxDb.Hsapiens.UCSC.hg19.knownGene

62. Hinrichs, Angela S., et al. “The UCSC genome browser database: update 2006.” Nucleic acids research 34.suppl_1 (2006): D590–D598. 10.1093/nar/gkj144

