## Supplemental Figures 1-8 for "Innate immune epigenomic landscape following controlled human influenza virus infection"

**Figure S1. Overview of RNA-seq and ATAC-seq fold-change correlations between naive HVL subjects and other (naive LVL and vaccinated HVL) subjects. (Related to Figure 2.)**

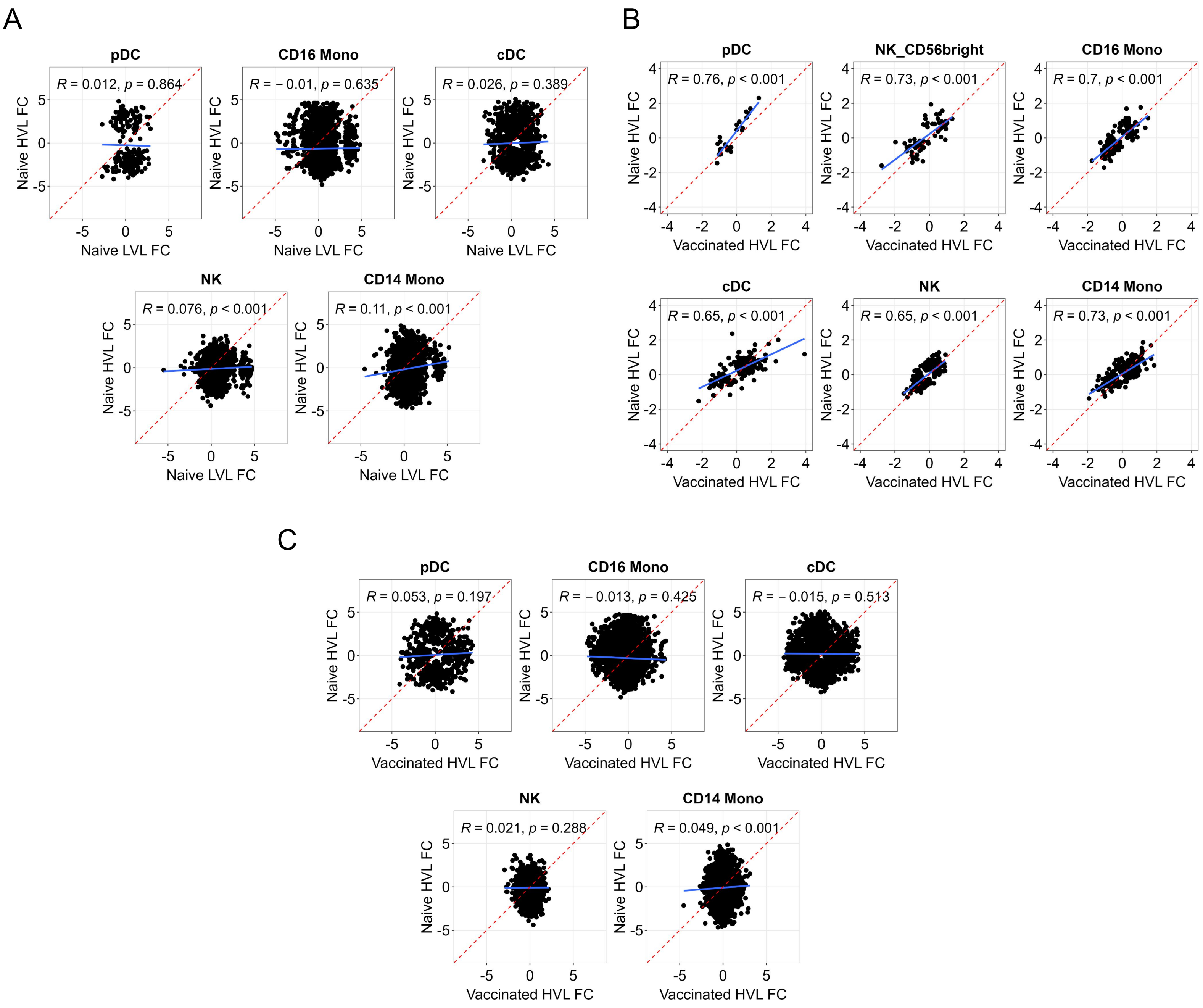

**Figure S2. Promoter sites that are downregulated at 28 days post-challenge are associated with inflammatory and active epigenetic remodeling processes. (Related to Figure 3.)**

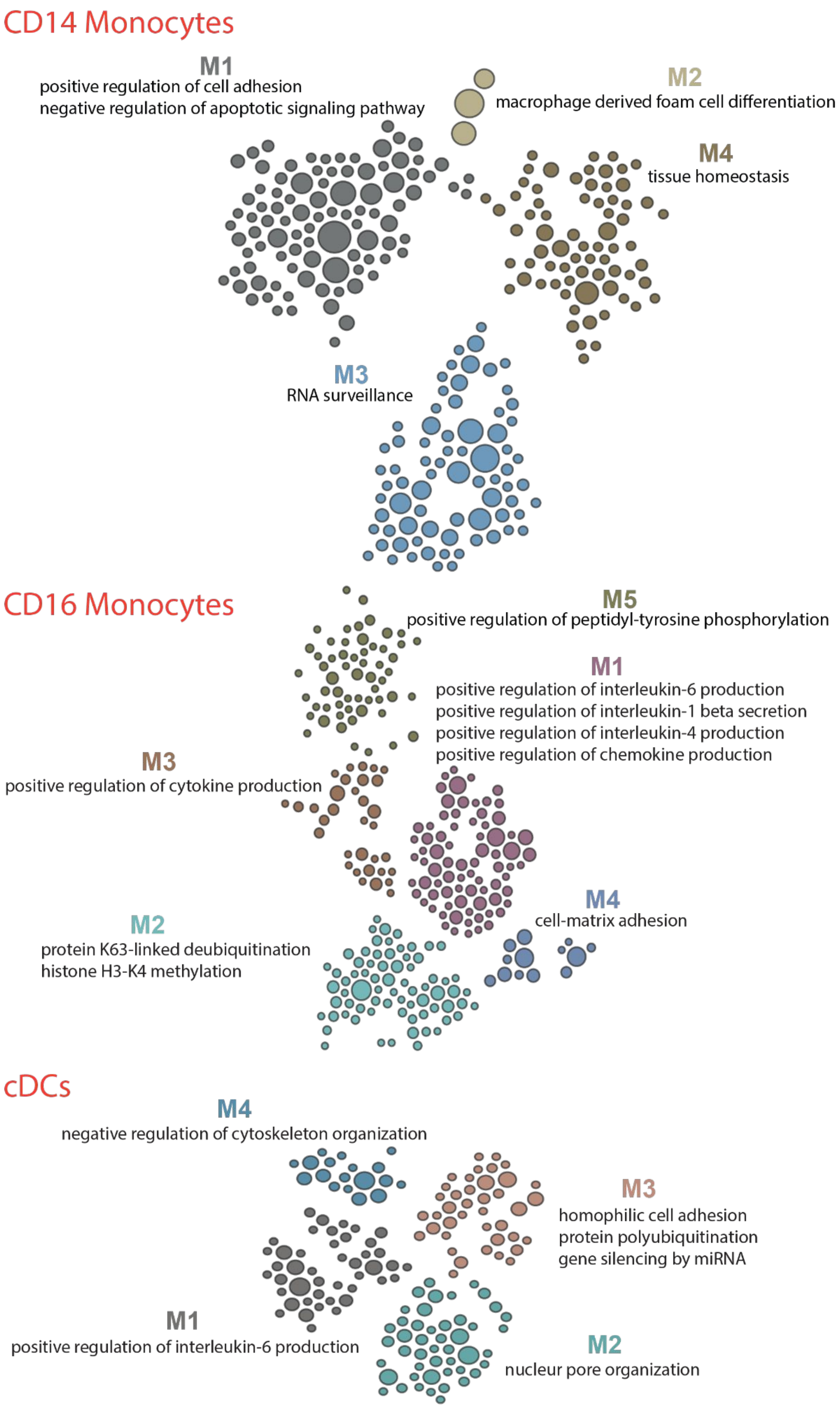

**Figure S3. Transcription factor motif enrichment analysis on differentially accessible sites from different innate immune cell types in naive LVL and vaccinated HVL subjects. (Related to Figure 4.)**

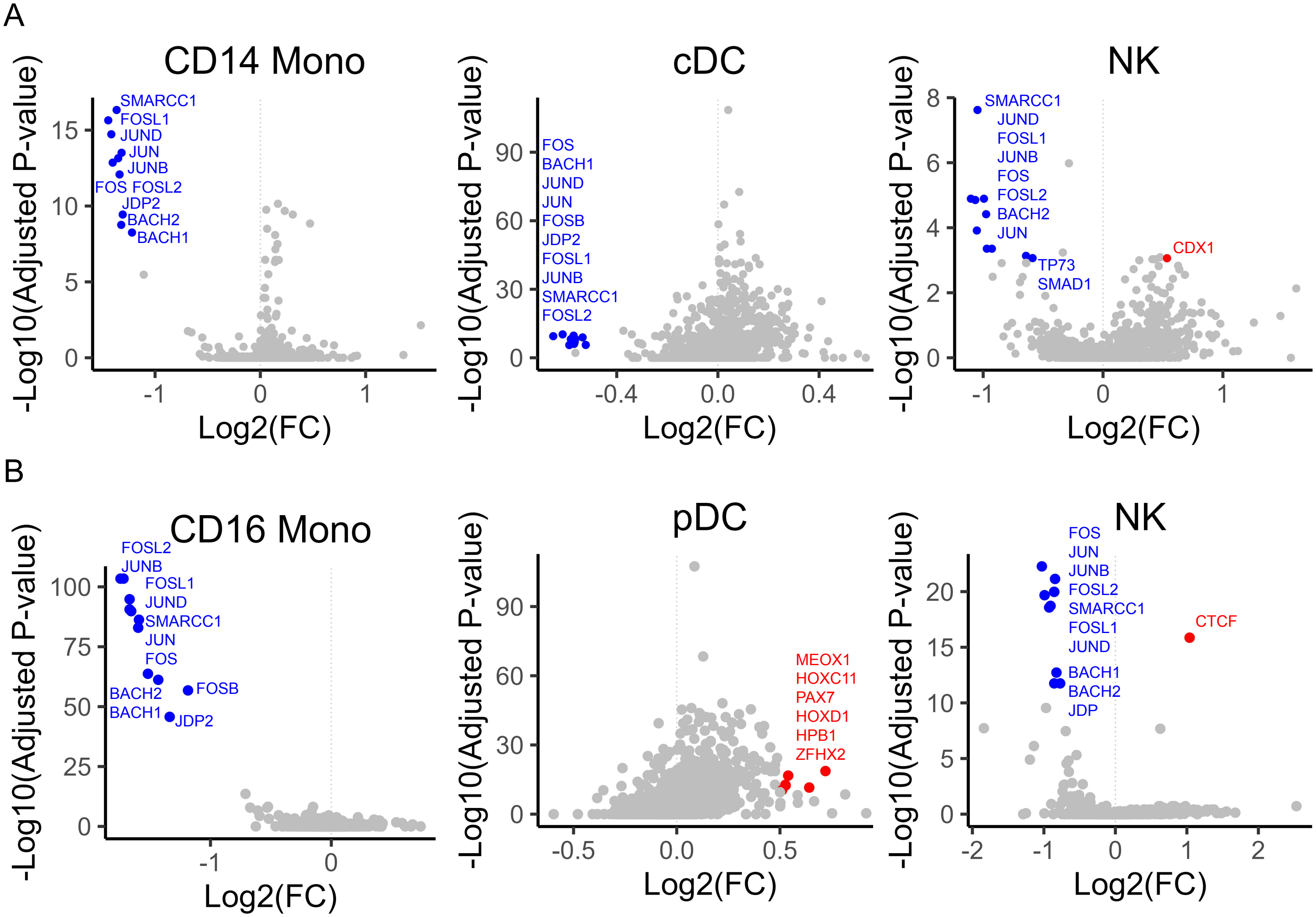

**Figure S4. Gene ontology enrichment analysis on upregulated DEG in CD14 monocytes. (Related to Figure 4.)**

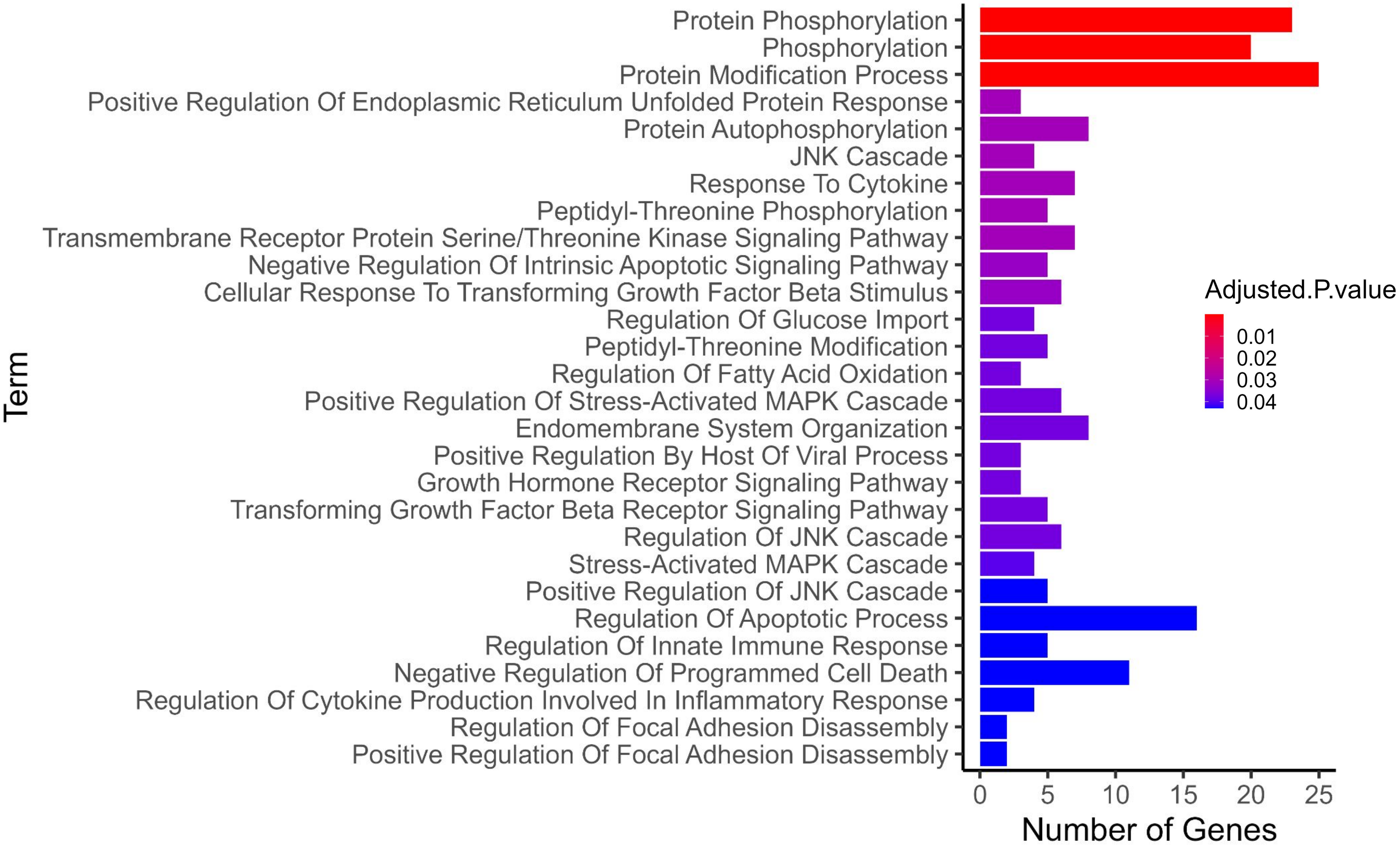

**Figure S5. Inflammatory-related DEG in vaccinated HVL and naive LVL subjects. (Related to Figure 4.)**

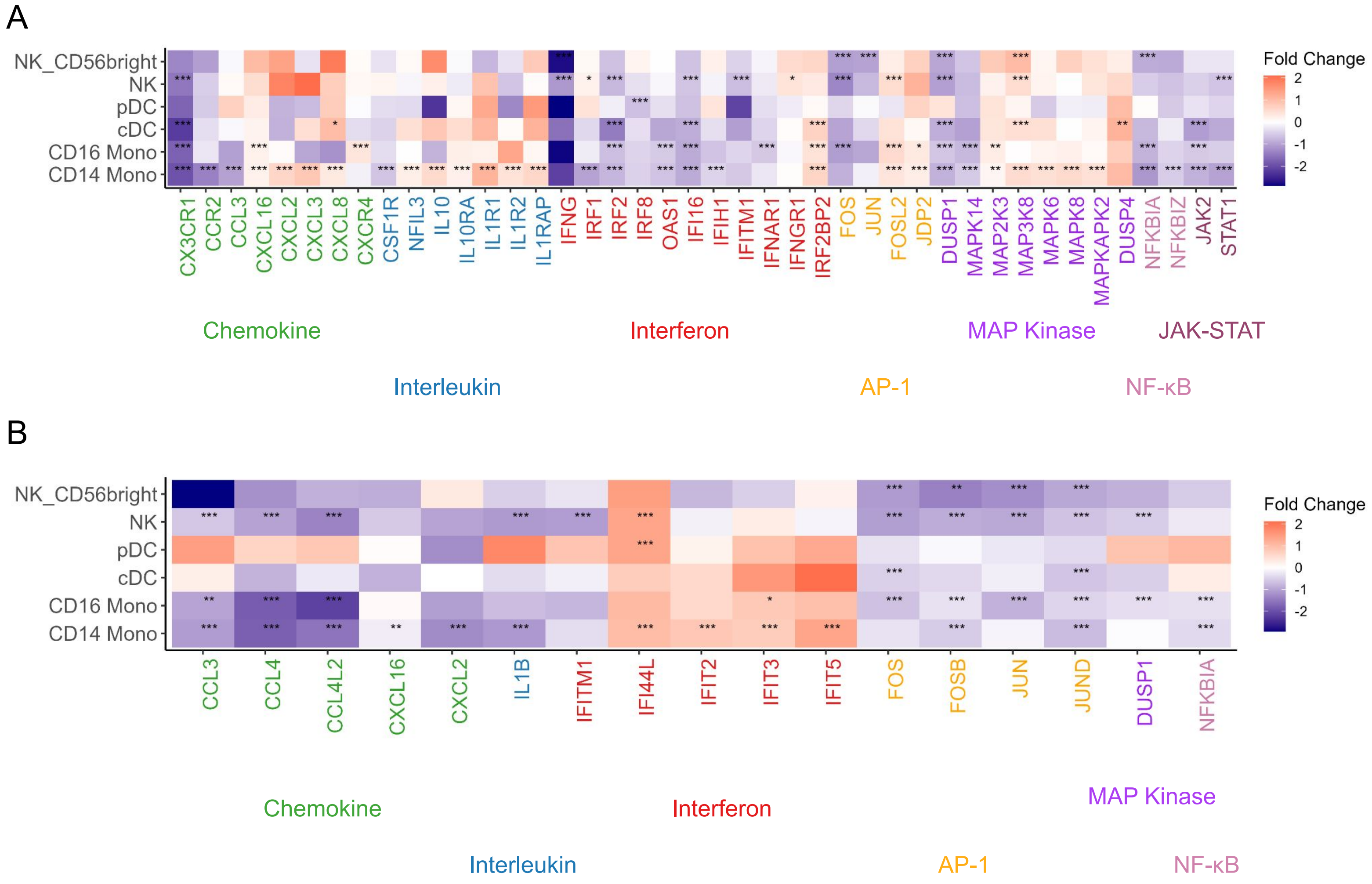

**Figure S6. Expression of inflammatory-related and epigenetic remodeling-related DEG across time course of infection. (Related to Figure 4.)**

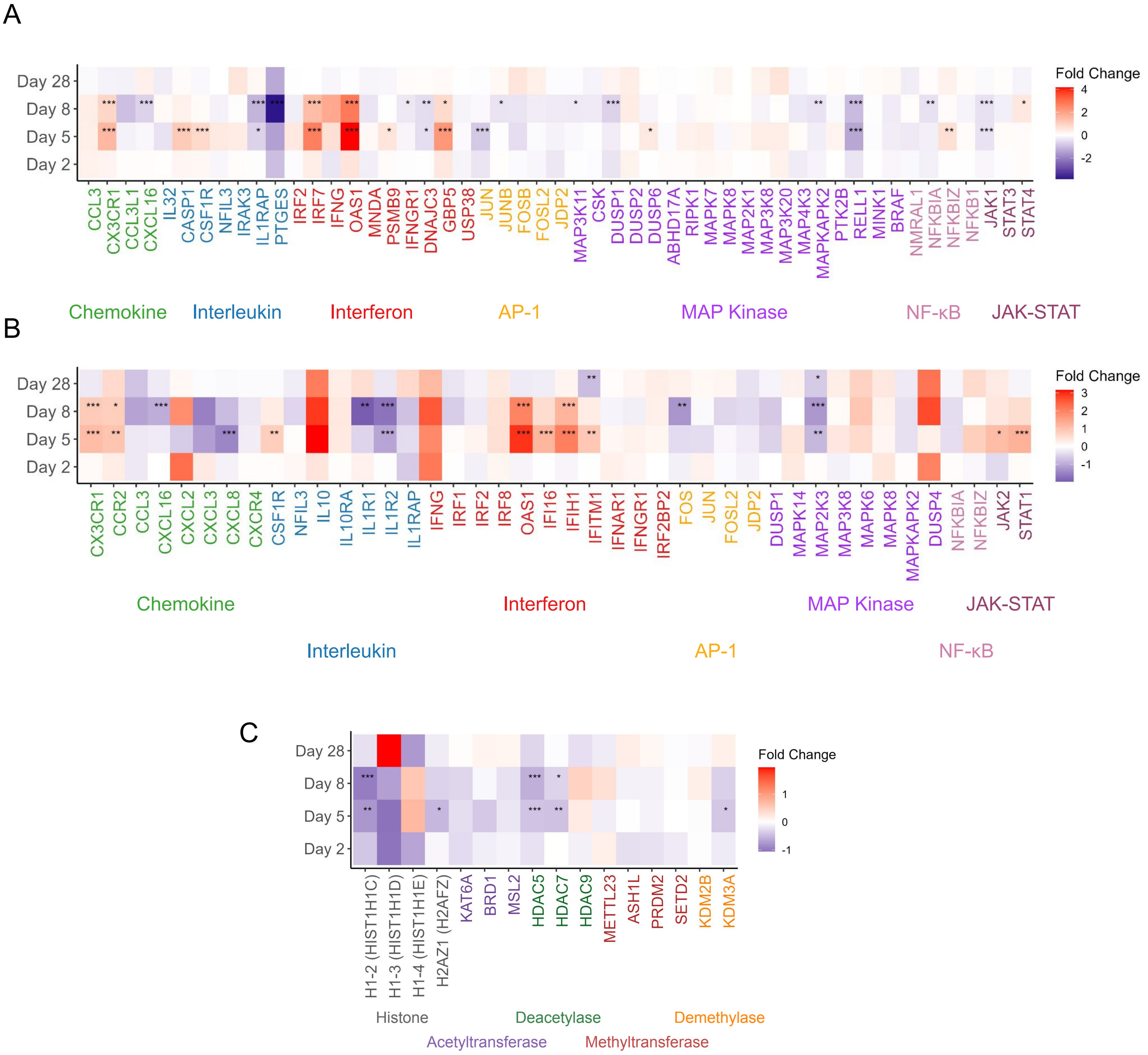

**Figure S7. Overview of fold change correlations between naive HVL subjects and other (naive LVL and vaccinated HVL) subjects for inflammatory-related and epigenetic-remodeling related genes. (Related to Figure 4.)**

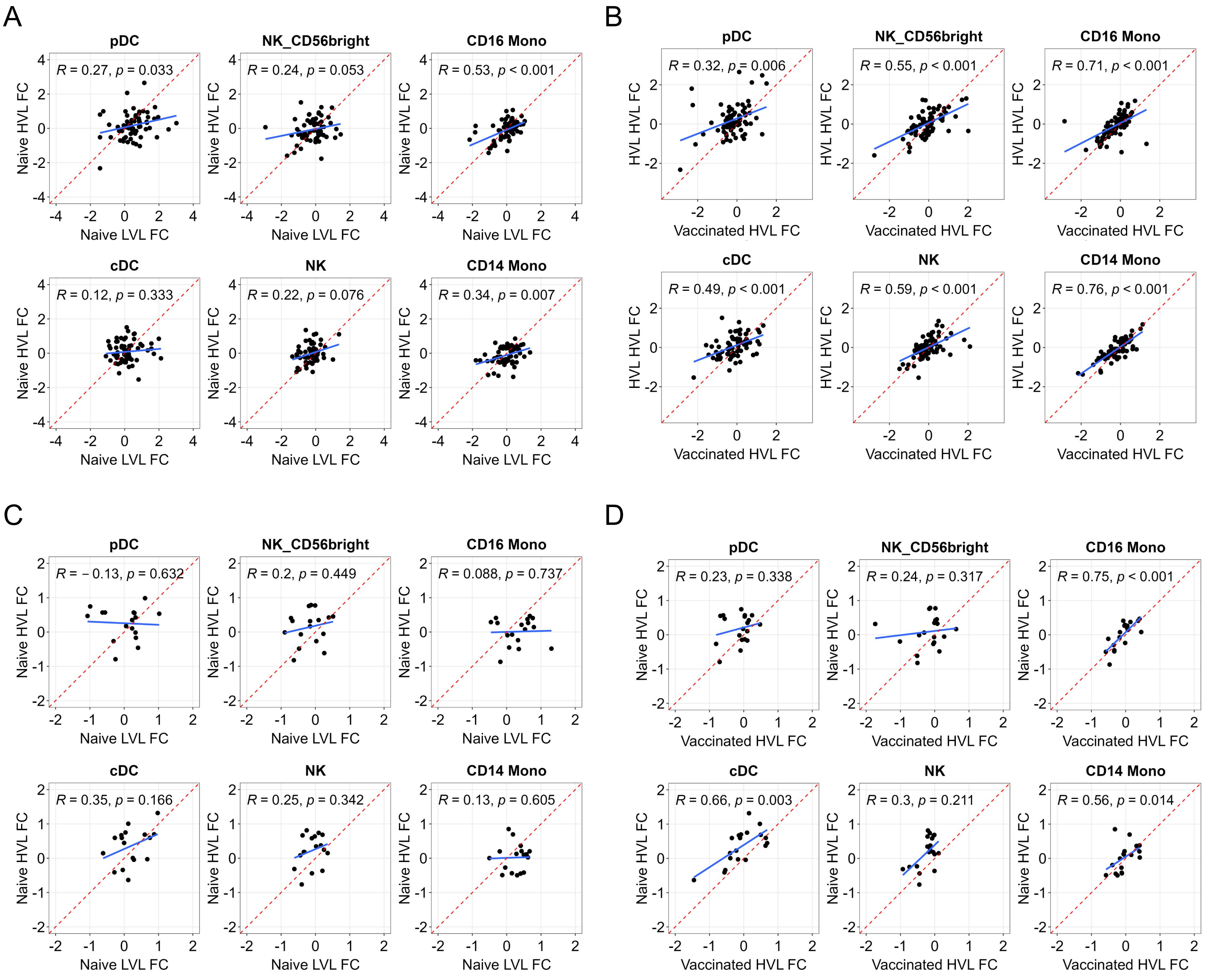

**Figure S8. Gene regulatory circuits in naive LVL and vaccinated HVL subjects share similar transcription factors to gene regulatory circuits in naive HVL subjects. (Related to Figure 5.)**

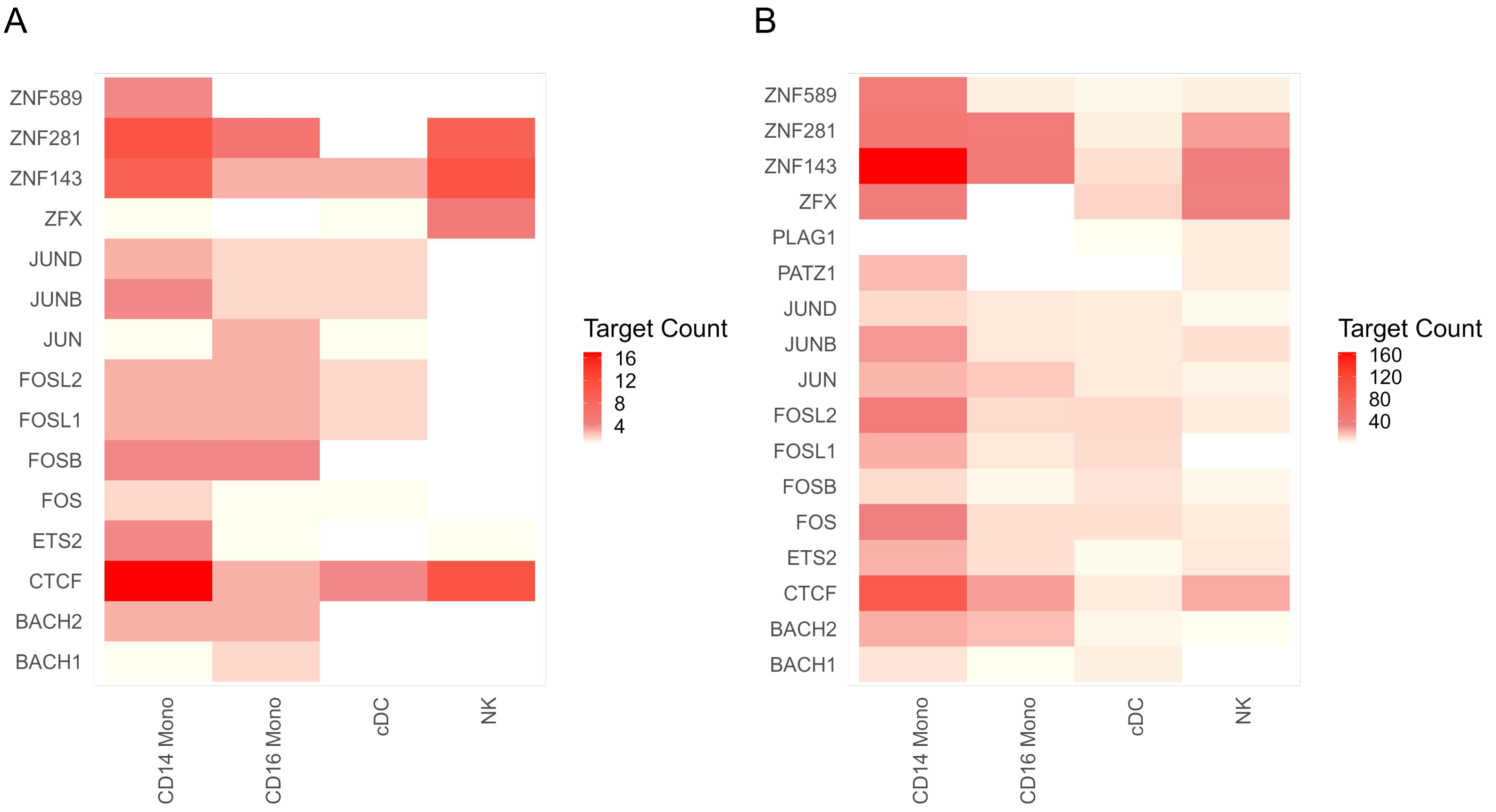
